# Ecophysiology of the cosmopolitan OM252 bacterioplankton (Gammaproteobacteria)

**DOI:** 10.1101/2021.03.09.434695

**Authors:** Emily R. Savoie, V. Celeste Lanclos, Michael W. Henson, Chuankai Cheng, Eric W. Getz, Shelby J. Barnes, Douglas E. LaRowe, Michael S. Rappé, J. Cameron Thrash

## Abstract

Among the thousands of species that comprise marine bacterioplankton communities, most remain functionally obscure. One key cosmopolitan group in this understudied majority is the OM252 clade of Gammaproteobacteria. Although frequently found in sequence data and even previously cultured, the diversity, metabolic potential, physiology, and distribution of this clade has not been thoroughly investigated. Here we examined these features of OM252 bacterioplankton using a newly isolated strain and genomes from publicly available databases. We demonstrated that this group constitutes a globally distributed novel genus (*Candidatus* Halomarinus), sister to *Litoricola*, comprising two subclades and multiple distinct species. OM252 organisms have small genomes (median 2.21 Mbp) and are predicted obligate aerobes capable of alternating between chemoorganoheterotrophic and chemolithotrophic growth using reduced sulfur compounds as electron donors, with subclade I genomes encoding the Calvin-Benson-Bassham cycle for carbon fixation. One representative strain of subclade I, LSUCC0096, had extensive halotolerance but a mesophilic temperature range for growth, with a maximum of 0.36 doublings/hr at 35°C. Cells were curved rod/spirillum-shaped, ~1.5 × 0.2 μm. Growth on thiosulfate as the sole electron donor under autotrophic conditions was roughly one third that of heterotrophic growth, even though calculations indicated similar Gibbs energies for both catabolisms. These phenotypic data show that some *Ca.* Halomarinus organisms can switch between serving as carbon sources or sinks and indicate the likely anabolic cost of lithoautotrophic growth. Our results thus provide new hypotheses about the roles of these organisms in global biogeochemical cycling of carbon and sulfur.

**Importance:** Marine microbial communities are teeming with understudied taxa due to the sheer numbers of species in any given sample of seawater. One group, the OM252 clade of Gammaproteobacteria, has been identified in gene surveys from myriad locations, and one isolated organism has even been genome sequenced (HIMB30). However, further study of these organisms has not occurred. Using another isolated representative (strain LSUCC0096) and publicly available genome sequences from metagenomic and single-cell genomic datasets, we examined the diversity within the OM252 clade, the distribution of these taxa in the world’s oceans, reconstructed the predicted metabolism of the group, and quantified growth dynamics in LSUCC0096. Our results generate new knowledge about the previously enigmatic OM252 clade and point towards the importance of facultative chemolithoautotrophy for supporting some clades of ostensibly “heterotrophic” taxa.

## Introduction

Marine bacterioplankton constitute 10^4^ to 10^7^ cells per milliliter in seawater (1–3), spread across hundreds to thousands of operational taxonomic units (OTUs) (2). However, many of these bacterioplankton lineages have no assigned metabolic or ecological roles and we know little more about them than their distribution in 16S rRNA gene surveys. While some of the dominant groups like SAR11 Alphaproteobacteria, *Prochlorococcus* Cyanobacteria, and SAR86 Gammaproteobacteria rightly attract considerable attention (4–7), many taxa that occur at somewhat lower relative abundances, but nevertheless are cosmopolitan microbial community members of the global oceans, have received comparably little study. One of these groups, the OM252 clade of Gammaproteobacteria, was first described over twenty years ago in clone library sequences from surface waters overlying the continental shelf off Cape Hatteras, North Carolina (8). This group is widely distributed. OM252 16S rRNA gene sequences have been reported from Sapelo Island off the coast of Georgia (9), the Gulf of Mexico (10–12), Kāneʻohe Bay in Oahu (13), the eutrophic coastal North Sea near Amsterdam (14), a lagoon in the Clipperton Atoll off the western coast of Mexico (15), and the Gulf of Lyon in the Mediterranean Sea (16). Sequences in GenBank with high percent identity to the OM252 clade also have come from near Cocos Island in the eastern tropical Pacific Ocean as well as the East China Sea. OM252 sequences occur in less saline waters, like the estuarine zone of the Jiulong River, China, and lakes with varying salinities in Tibet (17); but also hypersaline environments, like the Salton Sea in California (18), and salterns in Spain (19). There are even reports that indicate OM252 bacteria may be at least transiently associated with marine invertebrate microbiomes (20, 21). Thus, it appears that OM252 bacteria inhabit a variety of habitats and may have a euryhaline lifestyle.

Despite the widespread distribution of OM252 bacterioplankton, they remain poorly studied. The first reported isolate, HIMB30, was obtained via high-throughput dilution-to-extinction (DTE) cultivation with a natural seawater medium inoculated from Kāneʻohe Bay, Hawaiʻi (22). The ~2.17 Mbp HIMB30 genome predicted partial glycolysis, a complete TCA cycle, phototrophy via proteorhodopsin, carbon monoxide and sulfur oxidation, and CO_2_ fixation via the Calvin-Benson-Bassham cycle (22). However, these functions have not been demonstrated experimentally, nor have growth parameters, such as temperature or salinity tolerances, been investigated. We also don’t know how representative the HIMB30 features above are for the clade. Even the phylogenetic position of OM252 within the Gammaproteobacteria remains in question. The first clone library sequence branched sister to the OM182 clone and Oceanospirillales sequences (8). The closest described organisms, *Litoricola* spp., share less than 90% 16S rRNA gene identity with HIMB30 (22). Furthermore, the gammaproteobacterial phylogeny continues to evolve, with many traditionally recognized groups no longer remaining monophyletic (23), and additional genomes from uncultivated organisms changing the topology (24). The current Genome Taxonomy Database (GTDB release 05-RS95) indicates these organisms belong to the family Litoricolaceae, in the newly reconstituted Order Pseudomonadales (24–26).

Our previous work combining cultivation and cultivation-independent methods demonstrated that OM252 was a prominent member of the coastal northern Gulf of Mexico (10, 12). Our 16S rRNA gene amplicon data indicated at least two distinct amplicon sequence variants (ASVs) within the single observed OM252 OTU across six sampling sites and 3 different years (12). That OTU was the 25th most abundant bacterioplankton taxon in the high-salinity community (salinities > 12) observed in the three-year dataset. OM252 thus represented an important medium-abundance organism in that coastal environment. Furthermore, our artificial media facilitated ready cultivation of OM252 members, with over 30 strains isolated over the course of 17 experiments (12).

To improve our understanding of the physiology, ecology, and evolutionary relationships of the OM252 clade, we sequenced the genome of one representative isolate, LSUCC0096, and performed comparative genomic analyses with this organism, HIMB30, and 23 other publicly available environmental genomes. In parallel, we characterized physiological aspects of LSUCC0096 relevant to OM252 biology. OM252 comprised at least two subclades (I and II), both of which had a globally cosmopolitan distribution. OM252 clade members share many of the same metabolic features; however, there is subclade differentiation in the capacity for predicted sulfur-based chemolithoautotrophy. LSUCC0096 had a wide tolerance for salinity, growing from low salinity brackish water to nearly double the salinity of seawater. Furthermore, we showed that LSUCC0096 could grow under chemolithoautotrophic conditions with thiosulfate as the sole electron donor and estimate the energetic consequences of this metabolism on growth rates. Contrary to existing nomenclature in GTDB, our comparative genomic data support the designation of OM252 as a separate genus from *Litoricola*, which we propose as *Candidatus* Halomarinus, along with names for three species within the genus. These results expand our understanding of the genomic diversity, distribution, and lifestyles within the OM252 clade and provide the first cellular and physiological data for these organisms. They also raise new questions about the relationship between facultative chemolithotrophy and OM252 ecology.

## Materials and Methods

### LSUCC0096 isolation, genome sequencing, and genome assembly

Strain LSUCC0096 was isolated and initially identified via 16S rRNA gene PCR as previously reported (10) from surface water collected in Bay Pomme d’Or near the Mississippi River Birdfoot delta on January 12, 2015 (Buras, LA) (29.348784, −89.538171). DNA was extracted from cultures of LSUCC0096 that had reached max cell density (~10^6^ cells mL^−1^) growing in JW1 medium (10) at room temperature using a MoBio PowerWater DNA Isolation kit (QIAGEN, Massachusetts, USA) following the manufacturer’s protocols. Truseq DNA-seq Library preparation and Illumina MiSeq (paired-end 250 bp reads) sequencing was completed at the Argonne National Laboratory Environmental Sample Preparation and Sequencing Facility, producing 242,062 reads (Table S1). The genome was assembled using the A5 MiSeq pipeline (version 20150522) (27) with default settings. The LSUCC0096 genome was annotated at IMG (28) (Taxon ID 2639762503). For comparative genomics we re-annotated the genome along with other analyzed genomes using Anvi’o (see below), and the scaffolds were also deposited in GenBack (see Data Availability section).

### 16S rRNA gene phylogenies

The 16S rRNA gene of the LSUCC0096 genome was searched against both the NCBI nt and refseq_rna databases (accessed August, 2018) using megablast v. 2.2.28+ with –max_target_seqs 1000 and –num_threads 16. A selection of best hits was generated from each blast search and combined with the LSUCC0096 and HIMB30 16S rRNA genes from IMG, along with those from five different *Litoricola* spp. and the original OM252 clone library sequence U70703.1 (8). Additional 16S rRNA genes from the OM252 MAGs and SAGs were obtained from the Anvi’o genome database (see above) using the command anvi-get-sequences-for-hmm-hits --external-genomes external-genomes.txt -o 16S.fna --hmm-source Ribosomal_RNAs --gene Bacterial_16S_rRNA (or --gene Archaeal_16S_rRNA). The 16S rRNA gene of TOBG-NAT-109 had best blast (megablast online, default settings) hits to Bacteroides sequences, and was removed from further 16S rRNA gene analyses. The remaining sequences were aligned with MUSCLE v3.6 (29), culled with TrimAl v1.4.rev22 (30) using the -automated1 flag, and the final alignment was inferred with IQ-TREE v1.6.11 (31) with default settings and -bb 1000 for ultrafast bootstrapping (32). Tips were edited with the nw_rename script within Newick Utilities v1.6 (33) and trees were visualized with Archaeopteryx (34). Fasta files for these trees and the naming keys are provided at https://doi.org/10.6084/m9.figshare.14036573.

### Additional taxon selection

The HIMB30 genome (22) was downloaded from IMG (Taxon ID 2504557021). To provide a more comprehensive analysis of the OM252 clade beyond the LSUCC0096 and HIMB30 genomes, we searched for metagenome-assembled genomes (MAGs) that matched LSUCC0096 and HIMB30 using the following methods. We downloaded MAGs reconstructed from the Tara Oceans dataset (35, 36) and the northern Gulf of Mexico (37, 38). We identified all MAGs with average nucleotide identities (ANI) of > 76% to LSUCC0096 and HIMB30 using FastANI v1.1 (39) with default settings. These MAGs and the LSUCC0096 and HIMB30 genomes were then placed into the Genome Taxonomy Database (GTDB) tree [which also included additional MAGs constructed from the Tara Oceans dataset (40)] with GTDBtk v0.2.1 (24) (downloaded Feb 2019) using “classify_wf”. All genomes occurred in a monophyletic group including f_Litoricolaceae. The additional genomes from GTDB in this clade were downloaded. We then searched six representative genomes (LSUCC0096, HIMB30, GCA_002480175.1, GCA_002691485.1, UBA1114, UBA12265) against the Gammaproteobacteria Single-Amplified Genomes (SAGs) generated from GORG-Tropics collection (41) using FastANI as above. Finally, all genomes from this selection process were compared to each other with FastANI again. We then calculated the percent completion and contamination using CheckM v1.0.13 (42) using “lineage_wf”. We designated genomes as redundant if they had an ANI value of ≥ 99% with another genome. If a genome had a redundant match, we kept the genome with the highest % completion and lowest % contamination. Genomes with less than 50% estimated completion were discarded. The final genome selection statistics are in **Table S1**, available at https://doi.org/10.6084/m9.figshare.14067362.

### Phylogenomics

Based on the 16S rRNA gene phylogeny, we selected 208 genomes for a concatenated phylogenomic tree that spanned a variety of clades within the Gammaproteobacteria with members near OM252, plus 6 outgroup taxa from the Alpha- and Betaproteobacteria. These, together with 26 putative OM252 genomes (total 240), were analyzed using the Anvi’o phylogenomics pipeline through HMM assignment. Single copy marker genes that had membership in at least half the taxa in the tree (120) were selected using “anvi-get-sequences-for-hmm-hits --external-genomes external-genomes.txt -o temp.faa --hmm-source Rinke_et_al --return-best-hit --get-aa-sequences --min-num-bins-gene-occurs 120”, which returned a fasta file for each of the resulting 78 gene clusters. Each of these were aligned with MUSCLE v3.6 (29), culled with TrimAl v1.4.rev22 (30) using the -automated1 flag, and concatenated with the geneStitcher.py script from the Utensils package (https://github.com/ballesterus/Utensils) as described (43). The final alignment had 29,631 amino acid positions, and the tree was inferred with IQ-TREE v1.6.11 (31) with default settings and -bb 1000 for ultrafast bootstrapping (32). Tree tips were edited with the nw_rename script within Newick Utilities v1.6 (33) and trees were visualized with Archaeopteryx (34) and FigTree v1.4.3 and edited with Adobe Illustrator. The concatenated alignment and naming key is provided at https://doi.org/10.6084/m9.figshare.14036594.

### Pangenomics

Upon inspection of the phylogenomic tree, one putative genome, GCA_002408105.1, branched outside of the OM252 group (Fig. 1). We therefore excluded it from pangenomic analyses. The final 25 OM252 genomes were processed via Anvi’o v5.3 (44) using the pan-genomics workflow (45). This approach ensured that all genomes were subject to the same gene-calling and annotation workflow. Single copy marker genes via Anvi’o-provided HMMs and NCBI COGs were assigned, and Anvi’o-based gene calls were used for additional external annotation via Interproscan v5.33-72.0 (46) and KEGG assignments with GhostKOALA (47). All annotations are provided in Table S1 (https://doi.org/10.6084/m9.figshare.14067362) as part of the pangenome summary generated via Anvi’o: “anvi-summarize -p OM252/OM252pang-PAN.db -g OM252-GENOMES.db -C DEFAULT -o PAN_SUMMARY”. Metabolic reconstruction was completed using the KEGG annotations from GhostKOALA and a custom set of HMMs deployed with the KEGG-decoder, KEGG-expander, and Order_Decode_and_Expand scripts used previously (48). HMM searches for this workflow were completed using HMMER3.1b1 (49). Gene function enrichments based on annotations and pan-genome distribution were also calculated with Anvi’o. Using the phylogenomic clade structure (see below) that was also supported by ANI values, the subclades were imported into the Anvi’o database as layers using anvi-import-misc-data. Functional enrichments were then quantified for all the various annotation sources via the following command “for i in COG_CATEGORY Hamap ProSiteProfiles KeggGhostKoala SMART Gene3D TIGRFAM COG_FUNCTION SFLD PANTHER Coils CDD Pfam MobiDBLite ProSitePatterns PIRSF PRINTS SUPERFAMILY ProDom; do anvi-get-enriched-functions-per-pan-group -p OM252/OM252pang-PAN.db -g OM252-GENOMES.db --category subclade --annotation-source $i -o $i.enriched-subclade.txt; done”. ProgressiveMauve v2.4.0 (50) was used to align the HIMB30 and LSUCC0096 genomes using default settings and genbank files supplied from IMG. The OM252 Anvi’o pangenomic summary, including all annotations, is available in Table S1 (https://doi.org/10.6084/m9.figshare.14067362). The enriched function files are available at https://doi.org/10.6084/m9.figshare.14036579 and the ProgressiveMauve alignments are available https://doi.org/10.6084/m9.figshare.14036588.

**Figure 1.**
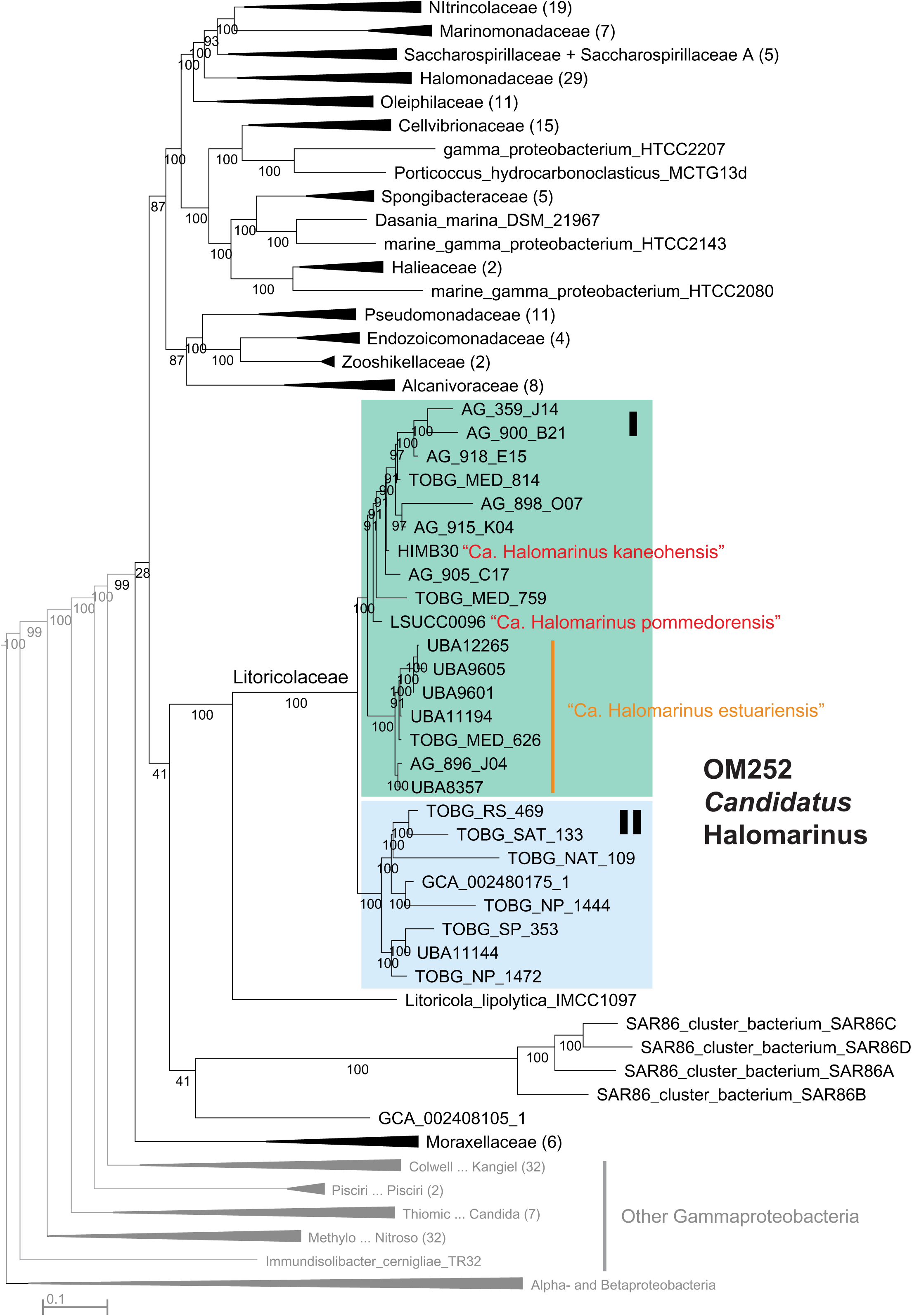
Phylogenomic tree of the Pseudomonadales and OM252. Maximum-likelihood tree based on 78 concatenated single-copy genes within the Pseudomonadales (as designated by GTDB) and selected other Gammaproteobacteria, with Alpha- and Betaproteobacteria outgroup. Final alignment = 29,631 amino acid positions. Families designated by GTDB within the Pseudomondales are indicated, with shading for the OM252 clade. Species designated in this study are highlighted in red and orange text. Values at nodes indicate bootstrap support (n=1000), scale indicated changes per position.

### Metagenomic read recruitment and analyses

Competitive recruitment of the metagenomic reads from Tara Oceans (2), BIOGEOTRACES (51), the Malaspina Global Expedition (52), the Southern California Bight near Los Angeles (53), the San Francisco Bay Estuary (with permission - C. A. Francis), and the northern Gulf of Mexico hypoxic zone (38) to the OM252 genomes was completed using the protocol available at http://merenlab.org/data/tara-oceans-mags/. Reads were cleaned using illumina utils v2.6 (54) implementing the method described in (55). Mapping used Bowtie 2 v2.3.2 (56), processing with SAMtools v0.1.19-44428cd (57), and read filtering with BamM v1.7.3 (http://ecogenomics.github.io/BamM/) to include only recruited hits with an identity of at least 95% and alignment length of at least 75%. The count table for each sample was generated using the get_count_table.py script from (https://github.com/edamame-course/Metagenome). Reads per kilobase per million (RPKM) calculations were performed using RPKM_heater (https://github.com/thrash-lab/rpkm_heater) and log_10_-transformed to improve visualization of recruitment across wide variations in abundance. RPKM calculations are available in Table S1 (https://doi.org/10.6084/m9.figshare.14067362). Visualization of the data for individual genome recruitment was completed in R (https://github.com/thrash-lab/metaG_plots). OM252 community diversity was assessed using the skbio.diversity algorithmic suite v0.5.6 (http://scikit-bio.org/docs/latest/diversity.html). Recruited OM252 reads from 588 metagenomic samples including TARA (2), BIOGEOTRACES (51), and Malaspina Global Expedition (52) were normalized to transcripts per million (TPM) values and used analogously for count dissimilarities. TPM calculations were performed using (https://github.com/thrash-lab/counts_to_tpm). The □-diversity algorithm (within http://scikit-bio.org/docs/latest/diversity.html) was retrofitted to interpret phylogenetic relationships using weighted-unifrac distances in place of the traditional Bray-Curtis dissimilarity. Retrofitting was performed *via* (https://github.com/thrash-lab/diversity_metrics). Sampling metadata for latitude, ocean region, depth, salinity, and temperature were collected to qualitatively assess the dissimilarity matrix in relation to designated intervals. ANOSIM correlation statistics were calculated for each metadata analysis treating absolute similarity of each OM252 community between samples as the null hypothesis to the alternative where community recruitment varies strongly with environmental metadata, such that: ANOSIM = 0 ≅ absolute similarity ≅ ubiquitous and even distribution across samples, and ANOSIM = 1 ≅ absolute dissimilarity ≅ highly varied distribution strongly related to an environmental factor (e.g., temperature). Three-dimensional ordination plots were constructed to visualize the principal coordinate analysis (PCoA) along the three foremost axes. The PCoA plots for latitude, ocean region, depth, salinity, and temperature are available in Fig. S6.

### RuBisCO phylogeny

The predicted large subunit for both the LSUCC0096 and HIMB30 RuBisCO genes were searched against ncbi nt via the web (March 2019). The top 100 hits from each were screened for redundant sequences and combined with the 8 additional near-full-length homologs in the OM252 group identified via Anvi’o (anvi-get-sequences-for-gene-clusters -p OM252/OM252pang-PAN.db -g OM252-GENOMES.db --gene-cluster-id GC_00001582 -o GC_00001582.faa) and a set of RuBisCO reference type genes (58). Genes were aligned with MUSCLE v3.6 (29), culled with TrimAl v1.4.rev22 (30) using the -automated1 flag, and the final alignment was inferred with IQ-TREE v1.6.11 (31) with default settings and -bb 1000 for ultrafast bootstrapping (32). The final tree was visualized with Archaeopteryx (34). The fasta file, script, and tree file are provided at https://doi.org/10.6084/m9.figshare.14036597.

### Growth experiments

Cell concentrations for all physiological experiments were measured via a Guava EasyCyte 5HT flow cytometer (Millipore, Massachusetts, USA) as previously reported (10, 59), and cells were grown in acid-washed polycarbonate flasks. The growth temperature range was tested in the isolation medium, JW1 (10), in triplicate, at 4°C, 12°C, 25°C, 30°C, 35°C, and 40 °C; using a refrigerator, Isotemp cooling incubator (Fisher), and benchtop heating incubators (Fisher) (all non-shaking). Salinity tolerance was tested using two different methods. First, we only altered the concentration of NaCl in the JW1 medium from 0-5%, producing a range of salinities from 8.66 to 63.5 (calculated from chlorinity according to S ‰ = 1.80655 Cl ‰ (60)). In the second method, we altered the concentration of all major ions proportionally, producing a salinity range of 0.36 to 34.8 as previously reported (61). In both approaches, all other media components (carbon, iron, phosphate, nitrogen, vitamins, and trace metals) were unaltered. All salinity growth experiments were conducted in triplicate and at room temperature.

We tested whether LSUCC0096 could grow under chemolithoautotrophic conditions by growing cells in base JW1 medium with 100 μM thiosulfate and all organic carbon sources excluded (aside from that possibly obtained via vitamins) for four consecutive growth cycles to eliminate the possibility of carryover from the seed culture grown in JW1. As a positive control, LSUCC0096 was grown in JW1 medium with the normal suite of carbon compounds (10). For a negative control, LSUCC0096 was grown in JW1 media with no added organic carbon aside from vitamins. During the fourth and final growth cycle, a second negative control was added where organic carbon and vitamins were both excluded. Cells for each condition were grown in triplicate. Cultures were counted once a day for the first two cycles to ensure transfers could be completed at the end of log phase. For the third and fourth growth cycles, cells were counted every 12 hours. The fourth growth cycle is depicted in Figure 5. Strain purity and identity were verified at the end of experiments using PCR of the 16S rRNA gene as previously reported (10). Growth rates for all experiments were calculated with the sparse-growth-curve script (https://github.com/thrash-lab/sparse-growth-curve). In brief, for each individual growth curve (cell density vs. time), the exponential phase is extracted and cell densities are transformed in natural log. Linear regression is performed on the ln(cell density) vs. time in the exponential phase. The specific growth rate is therefore the slope of the linear regression result. The doubling rate equals the specific growth rate divided by ln(2). The sparse-growth-curve script can do the exponential phase extractions automatically. It calculates the numerical differentiations (Δ ln(Cell density)/ Δdt) between two time points as the instantaneous growth rates. The different phases (lag, exponential, and stationary) are differentiated by performing decision tree regression. The exponential phase is the period with maximum orders of magnitudes converted to cell densities.

### Electron microscopy

To preserve cells for microscopy, we fixed 100mL of mid-exponential LSUCC0096 culture with 3% glutaraldehyde (Sigma Aldrich) and stored at 4°C overnight. We filtered the cells onto a 25mm diameter 0.2μm pore-sized isopore polycarbonate membrane filter (MilliporeSigma) via vacuum filtration and performed ethanol dehydration by soaking the filter for 25 minutes in each the following ethanol concentrations: 30%, 50%, 70%, 80%, 90%, 95%, and 100%. The filter was then put into a Tousimis 815 critical point dryer and sputter coated for 45s in a Cressington 108 Manual Sputter Coater. Cells were imaged using the JSM-7001F-LV scanning electron microscope at the University of Southern California Core Center of Excellence in Nano Imaging (http://cemma.usc.edu/) with a working distance of 6.8mm and 15.0kV.

### Energetics

Overall Gibbs energies, Δ*G_r_*, of thiosulfate and organic carbon oxidation,

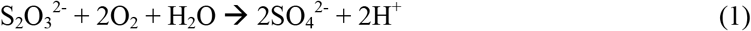

and

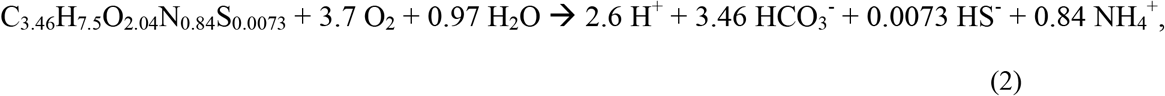

were calculated using

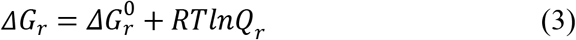

where 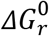 and *Q_r_* refer to the standard molal Gibbs energy and the reaction quotient of the indicated reaction, respectively, *R* represents the gas constant, and *T* denotes temperature in Kelvin. Values of 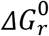 were calculated using the revised-HKF equations of state (62–64), the SUPCRT92 software package (65), and thermodynamic data taken from a number of sources (66–68). Values of *Q_r_* are calculated using

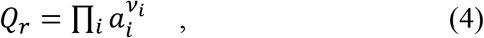

where *a_i_* stands for the activity of the *i*th species and *v_i_* corresponds to the stoichiometric coefficient of the *i*th species in the reaction of interest. Because standard states in thermodynamics specify a composition (69, 70) values of *Q_r_* must be calculated to take into account how environmental conditions impact overall Gibbs energies. In this study we use the classical chemical-thermodynamic standard state in which the activities of pure liquids are taken to be 1 as are those for aqueous species in a hypothetical 1 molal solution referenced to infinite dilution at any temperature or pressure.

Activities are related to concentration, *C*, by

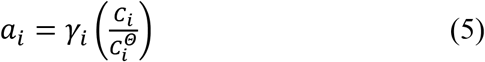

where *γ_i_* and *C_i_* stand for the individual activity coefficient and concentration of the *i*th species, respectively, and 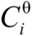 refers to the concentration of the *i*th species under standard state conditions, which is taken to be equal to one molal referenced to infinite dilution. Values of *γ_i_* were computed using an extended version of the Debye-Hückel equation (71)). Concentration of the species shown in Reactions (1) and (2) are those used in the media ([O_2_] = saturation in seawater at 25°C (205 μmol); [C_org_] = 66.6 μM; [HCO_3_^−^] = seawater (2 mmol); [HS^−^] = oxic seawater (0.5 nM)).

Because it is not clear which organic compounds are being oxidized for energy, we calculated values of Δ*G_r_* for this process by representing DOC as a weighted average of all of the organic compounds in the media recipe, shown on the left side of reaction (2). This composite formula was used to calculate the standard state Gibbs energy of reaction (2), 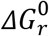, according to the algorithm given in LaRowe and Van Cappellen (2011), which relates the nominal oxidation of carbon, NOSC, in organics to their Gibbs energy of oxidation (the average weighted NOSC in the media is −0.26).

## Results

### Isolation and genome sequencing

LSUCC0096 was isolated as part of a series of DTE experiments using water samples from across the southern Louisiana coast (10). The specific sample from which we obtained LSUCC0096 came from surface water in the Bay Pomme d’Or near the Mississippi River Birdfoot delta; salinity 26, 7.7°C, pH 7.99. LSUCC0096 was grown for genome sequencing in JW1 medium (10). Illumina MiSeq PE 250bp sequencing generated 242,062 reads. Assembly with the A5 MiSeq pipeline resulted in 4 scaffolds with a total length of 1,935,310 bp, N50 of 1,442,657 bp, 30x median coverage, and a GC content of 48.5% (Table 1). Annotation by IMG predicted 2,001 protein-coding genes and 46 RNA genes- one copy of the 5S, 16S, and 23S rRNA genes and 36 predicted tRNA genes. The genome was estimated to be 96.17% complete with 0.37% contamination and a coding density of 95% (via CheckM (42), Table 1).

**Table 1.** Genome characteristics.

| Bin | SC | Est. Compln (%) | Est. Contam (%) | Strain het (%) | Genome size (bp) | Est. genome size (bp) | N50 scaffolds | Scaffolds | Longest scaffold | GC fract. | Coding density fract. | Type | Source | Study |
| --- | --- | --- | --- | --- | --- | --- | --- | --- | --- | --- | --- | --- | --- | --- |
| AG-359-J14 | I | 63.37 | 0 | 0 | 1319993 | 2082994 | 41035 | 55 | 132304 | 0.486 | 0.918 | SAG | BATS | Pachiadaki et al. 2019 |
| AG-900-B21 | I | 53.83 | 0 | 0 | 1259049 | 2338936 | 36080 | 65 | 106558 | 0.486 | 0.909 | SAG | BATS | Pachiadaki et al. 2019 |
| AG-918-E15 | I | 71.08 | 0.37 | 0 | 1671916 | 2352161 | 62227 | 52 | 215445 | 0.488 | 0.927 | SAG | BATS | Pachiadaki et al. 2019 |
| TOBG-MED-814 | I | 90.2 | 6.51 | 84 | 1967603 | 2181378 | 39535 | 74 | 144762 | 0.498 | 0.937 | MAG | Mediterranean Sea | Tully et al. 2018 |
| AG-898-O07 | I | 56.73 | 0 | 0 | 1067362 | 1881477 | 29177 | 60 | 110676 | 0.486 | 0.917 | SAG | BATS | Pachiadaki et al. 2019 |
| AG-915-K04 | I | 51.64 | 0 | 0 | 1290806 | 2499624 | 53286 | 58 | 157605 | 0.486 | 0.920 | SAG | BATS | Pachiadaki et al. 2019 |
| HIMB30 | I | 95.99 | 0.49 | 0 | 2168870 | 2259475 | 223986 | 36 | 638152 | 0.499 | 0.941 | Isolate | Kāne'ohe Bay, HI | This study |
| AG-905-C17 | I | 68.02 | 0 | 0 | 1505956 | 2213990 | 89387 | 53 | 220750 | 0.483 | 0.911 | SAG | BATS | Pachiadaki et al. 2019 |
| TOBG-MED-759 | I | 89.74 | 3.21 | 93.33 | 1788781 | 1993293 | 84972 | 31 | 211681 | 0.506 | 0.940 | MAG | Mediterranean Sea | Tully et al. 2018 |
| LSUCC0096 | I | 96.17 | 0.37 | 0 | 1935310 | 2012384 | 1442657 | 4 | 1442657 | 0.487 | 0.952 | Isolate | Bay Pomme d'Or, LA | This study |
| UBA12265 | I | 70.95 | 1.91 | 25 | 1444999 | 2036644 | 8081 | 220 | 28363 | 0.510 | 0.856 | MAG | Adriatic Sea | Parks et al. 2017 |
| UBA9605 | I | 69.58 | 1.7 | 11.11 | 1221408 | 1755401 | 6540 | 213 | 25083 | 0.511 | 0.912 | MAG | Inner Oslofjord | Parks et al. 2017 |
| UBA9601 | I | 77.59 | 0 | 0 | 1349581 | 1739375 | 8473 | 186 | 32481 | 0.511 | 0.913 | MAG | Outer Oslofjord | Parks et al. 2017 |
| UBA11194 | I | 68.83 | 0.75 | 0 | 1212584 | 1761709 | 6292 | 216 | 24783 | 0.511 | 0.916 | MAG | Adriatic Sea | Parks et al. 2017 |
| TOBG-MED-626 | I | 80.2 | 2.1 | 73.33 | 1193441 | 1488081 | 23640 | 59 | 71869 | 0.509 | 0.958 | MAG | Mediterranean Sea | Tully et al. 2018 |
| AG-896-J04 | I | 70.54 | 0.46 | 0 | 1622917 | 2300705 | 96547 | 37 | 166779 | 0.492 | 0.923 | SAG | BATS | Pachiadaki et al. 2019 |
| UBA8357 | I | 65.76 | 1.23 | 40 | 1312486 | 1995873 | 8992 | 180 | 28717 | 0.510 | 0.823 | MAG | Sapelo Island, GA | Parks et al. 2017 |
| TOBG-RS-469 | II | 69.27 | 2.04 | 16.67 | 1799083 | 2597204 | 14438 | 138 | 66997 | 0.494 | 0.932 | MAG | Red Sea | Tully et al. 2018 |
| TOBG-SAT-133 | II | 75.37 | 1.19 | 0 | 1870654 | 2481961 | 27755 | 73 | 67858 | 0.491 | 0.929 | MAG | South Atlantic | Tully et al. 2018 |
| TOBG-NAT-109 | II | 85.36 | 2.9 | 7.69 | 2285316 | 2677268 | 25595 | 107 | 89471 | 0.487 | 0.925 | MAG | North Atlantic | Tully et al. 2018 |
| GCA_002480175.1 | II | 83.79 | 2.75 | 27.27 | 2207349 | 2634382 | 10539 | 273 | 37539 | 0.485 | 0.851 | MAG | South Western Atlantic | Parks et al. 2017 |
| TOBG-NP-1444 | II | 83.86 | 3.15 | 11.11 | 2099531 | 2503614 | 26349 | 109 | 70809 | 0.475 | 0.911 | MAG | North Pacific | Tully et al. 2018 |
| TOBG-SP-353 | II | 53.45 | 0 | 0 | 1212569 | 2268604 | 23702 | 51 | 50076 | 0.482 | 0.924 | MAG | South Pacific | Tully et al. 2018 |
| UBA11144 | II | 72.06 | 1.95 | 20 | 1667084 | 2313467 | 8361 | 246 | 24773 | 0.483 | 0.857 | MAG | South Eastern Pacific | Parks et al. 2017 |
| TOBG-NP-1472 | II | 76.68 | 1.79 | 0 | 1521562 | 1984301 | 19361 | 94 | 46684 | 0.488 | 0.929 | MAG | North Pacific | Tully et al. 2018 |

Genomes shaded according to subclade (SC)

### Taxonomy

Initial blast searches of the 16S rRNA gene sequence to GenBank identified LSUCC0096 as a gammaproteobacterium, with the closest cultivated representative being the OM252 clade organism HIMB30 (22). To better understand the phylogenetic breadth of this group we identified 23 non-redundant good or high-quality MAGs and SAGs closely related to HIMB30 and/or LSUCC0096 based on average nucleotide identity (ANI) and monophyletic grouping within the Genome Taxonomy Database (GTDB) (Table S1). Phylogenetic inference using 16S rRNA gene sequence phylogenies produced different results depending on taxon selection (Figs. S1 and S2). OM252 clade sequences branched sister to the *Litoricola* genus in the RefSeq tree (Fig. S1), but with substantial evolutionary distance between them. However, with added diversity contributed by clones and other non-RefSeq sequences, this relationship did not hold (Fig. S2). *Litoricola* branched in a completely different part of the tree, whereas the OM252 clone library sequence (U70703.1) remained in a monophyletic group containing the genomes in this study (Fig. S2).

To improve the placement of the OM252 clade within the Gammaproteobacteria and test the sister relationship with *Litoricola*, we created a phylogenomic tree using concatenated single copy marker genes from OM252 and other Gammaproteobacteria genomes selected based on the 16S rRNA gene trees. Consistent with the RefSeq 16S rRNA gene tree, the OM252 clade branched sister to *Litoricola*, which together were sister to the SAR86 clade (Figs. 1, S3). This group branched between the Moraxellaceae and the remainder of the newly recircumscribed Pseudomonadales Order in GTDB (Figs 1, S3). Even though the tree contains all the major families designated by GTDB in the Pseudomonadales, the relationship of the Litoricolaceae does not match their current topology (https://gtdb.ecogenomic.org/, accessed February 2021), which places Litoricolaceae sister to the Saccharospirillaceae and Oleiphilaceae. However, the bootstrap support for the relationships with SAR86 and the branch leading to the rest of the Pseudomonadales was poor (Figs. 1, S3). Subclade structure within the OM252 clade (subclades I and II, Fig. 1) corresponded to subgroups circumscribed via FastANI during our taxon selection (Table S1). Within subclade I, a monophyletic group of genomes (UBA12265 to UBA8357 - Fig. 1) represented a single species according to the 95% ANI species cutoff (39), with multiple additional species in both subclades (Table S1). The isolates HIMB30 and LSUCC0096 shared only 80.3% ANI, making them distinct species.

Pairwise blast of the 16S rRNA gene from all nine OM252 subclade I members for which the gene was recovered with five *Litoricola* representatives from multiple species (Figs. S1, S2) corroborated the phylogenetic separation of these groups: no *Litoricola* sequence had greater than 89.8% identity with any OM252 genome, whereas the range of identity within OM252 subclade I was ≥ 98.5% (Table S1; no legitimate 16S rRNA genes were recovered from subclade II). Thus, OM252 subclade I constitutes a distinct genus from *Litoricola* based on pairwise 16S rRNA gene identity alone (72). The monophyletic relationship of subclade I and II in the phylogenomic tree, the presence of multiple distinct species within both subclades based on ANI, as well as the comparative branch length distances between subclades I and II vs. *Litoricola*, support inclusion of subclade I and II into the same group. Finally, there is a considerable difference in the GC content of all OM252 genomes (both subclades) compared with *Litoricola* (47-51% vs. 58-60% (26, 73), respectively). Thus, we propose the provisional genus name *Candidatus* Halomarinus for the OM252 clade. Since HIMB30 was the first reported isolate from OM252, this would be the type strain. However, since it is not currently deposited in international culture collections, we propose the species name as *Candidatus* Halomarinus kaneohensis, sp. nov.. We also propose *Candidatus* Halomarinus pommedorensis, sp. nov., for strain LSUCC0096, and *Candidatus* Halomarinus estuariensis, sp. nov., for the species cluster comprising UBA12265 to UBA8357 in Figure 1. We provide genus and species descriptions below.

### Distribution

We previously reported the distribution of the OM252 clade within the 16S rRNA gene amplicon data associated with three years of sampling in support of DTE experiments from the Louisiana coast (12). The single OM252 OTU was moderately abundant (relative abundance up to ~1%) at salinities > 5, regardless of site. We also identified two amplicon sequence variants (ASVs) associated with the OM252 clade- one which was generally much more abundant than the other. LSUCC0096 matched the more abundant, and more frequently cultivated, ASV5512 (12), representative of subclade I. ASV5512 was found across a range of salinities, but was more prevalent in salinities above 12, where it was one of the top 50 most abundant ASVs in the three-year dataset.

We expanded our assessment of OM252 genome abundance and distribution using metagenomic read recruitment for the global oceans. OM252 members from both subclades recruited reads from metagenomic samples across the globe (Fig. S4). The two most abundant taxa were represented by the subclade II MAGs TOBG-NAT-109 and GCA_002480175 (Fig. S5). The two most abundant subclade I taxa were represented by the SAGs AG-905-C17 and AG-900-B21. Genomes from the isolate strains LSUCC0096 and HIMB30 were the third and sixth ranking by median recruitment values for subclade I. Recruitment to the *Ca.* H. estuariensis cluster genomes was generally lower than to other genomes in subclade I (Fig. S5). Assessment of recruitment-based abundance patterns to the entire OM252 clade revealed no strong relationships with latitude, salinity, temperature, region, or depth (Fig. S6).

Individual genome recruitment was not influenced by salinity, but the salinity variation in the tested samples was quite limited (Fig. S7). Some genomes showed trends consistent with recruitment based on temperature (Fig. S8). For example, HIMB30 and LSUCC0096 RPKMs had significant negative relationships with temperature (linear regression, P-values 0.00168 and 0.00289, respectively). However, there was no consistent pattern for the genomes within a given subclade, and many genomes had no significant recruitment relationship with temperature (Fig. S8). The vast majority of samples from the dataset were from the epipelagic, and recruitment to OM252 genomes predominated in surface waters (Fig. S9). We observed very high relative recruitment to HIMB30 and AG-898-O07 in bathy- and abyssopelagic waters, and intermediate relative recruitment in deep ocean samples to other genomes from both subclades (Fig. S9), suggesting that some strains of OM252 may either preferentially or transiently inhabit the ocean interior. We also examined latitudinal distribution in recruitment. The dataset had a bimodal distribution with the majority of sites occurring in the mid-latitudes (Fig. S10). Most genomes did not show recruitment patterns consistent with the sample distribution alone, nor did we observe any clear relationships between subclade genomes with latitude (Fig. S11).

Separately, we recruited metagenomes from two coastal and one estuarine site- namely, the northern Gulf of Mexico “dead zone” (38); samples from the San Pedro shelf, basin and Catalina Island in the Southern California Bight (53); and a transect of samples from the San Francisco Bay estuary (Fig. S12). In contrast to the recruitment in the global oceans, the *Ca.* H. estuariensis cluster of genomes were the most abundant across these coastal/estuarine sites (Fig. S13), and in particular, were the highest recruiting members in the most saline sites of the San Francisco Bay estuary (Fig. S14). Subclade II was nearly absent in that same system. Thus, we hypothesize that the *Ca.* H. estuariensis group represents an estuarine-adapted species within OM252. Within the coastal/estuarine dataset, subclade I members generally showed increasing relative abundance with decreasing salinities, whereas subclade II members showed the opposite trends, although the data was sparse (Fig. S15).

### General genome characteristics

The shared and variable gene content and corresponding metabolic functions of the OM252 genomes are shown in Table 1. Estimated genome completion spanned 51.6-96.2%, with LSUCC0096 the most complete. Median estimated complete genome size was 2.21 Mbp, median GC content was 49% (47-51%), and median coding density was 92% (82-96%) (Table S1). Genome sizes are comparable to the *Litoricola lipolytica* IMCC1097 genome, the nearest phylogenomic neighbor. However, the IMCC1097 genome has a much higher GC content (58.8%) than that of *Ca.* Halomarinus.

### Electron transport and energy conservation

*Ca.* Halomarinus bacteria were predicted to be aerobic chemotrophs (Fig. 2, Table S1). No alternative terminal electron accepting processes were identified in either subclade (Fig. S16). A second, high-affinity (*cbb_3_*-type) cytochrome c oxidase was additionally present in seven subclade I genomes (Fig. 2, Table S1). Both subclades had sodium-translocating respiratory NADH dehydrogenases and Na+/H+ F-type ATPases, indicating the likely use of a sodium-motive force (Fig. 2). Most *Ca.* Halomarinus genomes contained proteorhodopsin (18/25) and retinal biosynthesis (19/25), with one notable exception being LSUCC0096. The proteorhodopsin gene in HIMB30 is located in an indel region with neighboring sections syntenic to the second largest contig in the LSUCC0096 genome (see ProgressiveMauve alignment in Supplementary Information). Therefore, it appears that the gene is truly absent from LSUCC0096 rather than missing as a result of the genome being incomplete.

**Figure 2.**
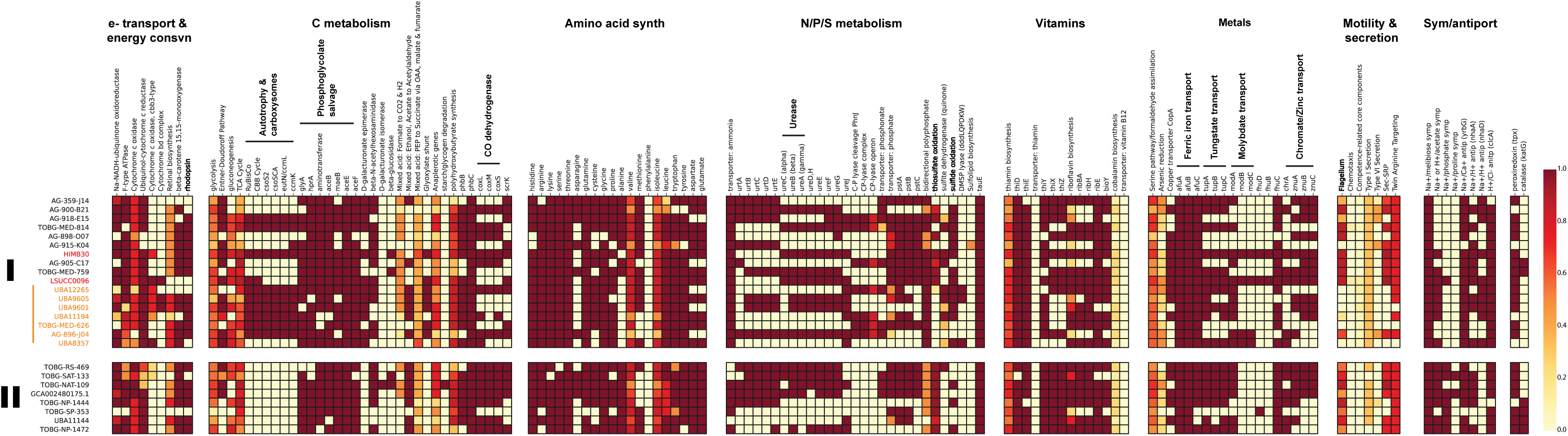
Metabolic reconstruction of the OM252 clade. Heatmap displays gene and pathway content according to the scale on the right. Subgroups of processes and key metabolic pathways are highlighted for ease of viewing. Subclades and species designations follow that in Figure 1.

### Carbon

Both subclades had predicted genes for the Entner-Doudoroff pathway, the TCA cycle, gluconeogenesis, most of the genes of the pentose-phosphate pathway, and fructokinase (*scrK*) for fructose utilization (Fig. 2, Table S1). Six subclade I genomes, including HIMB30 and LSUCC0096, had an annotated mannose-6-phosphate isomerase for mannose utilization. No genome had an annotated phosphofructokinase gene for glycolysis through the Embden-Meyerhof-Parnas pathway. The *coxMSL* aerobic carbon monoxide dehydrogenase genes originally reported in HIMB30 (22) were conserved among most genomes (e.g., *coxL* in 16/25) in *Ca.* Halomarinus, except LSUCC0096 (Table S1). Similar to proteorhodopsin, this deletion in LSUCC0096 occured with flanking regions of conservation to the HIMB30 genome and were not near any contig boundaries, making it likely this is a true gene deletion in the LSUCC0096 genome. Subclade I, but not II, had predicted genes for the glyoxylate bypass. Eighteen genomes also contained a predicted beta-N-acetylhexosaminidase (Fig. 2), a glycoside hydrolase of the CAZyme GH-20 family that may confer chitin-degradation capabilities on *Ca.* Halomarinus bacteria (74, 75).

Ten of the *Ca.* Halomarinus genomes in subclade I had a ribulose-1,5-bisphosphate carboxylase (RuBisCO) gene and the associated Calvin-Benson-Bassham pathway for carbon fixation (Fig. 2), making these organisms predicted facultative autotrophs. Phylogenetic analysis of the large RuBisCO subunit demonstrated that all were Type I RuBisCO genes; however, the LSUCC0096 large subunit grouped away from that of HIMB30 and the other *Ca.* Halomarinus genomes for which a sequence was recovered (Fig. S17). The LSUCC0096 RuBisCO genes were located directly upstream of likely *cbbQ* and *cbbO* activase genes (76), whereas the HIMB30 RuBisCO genes were located upstream from a suite of alpha-carboxysome genes (*csoS2, csoSCA, ccmL, ccmK*). Although we found carboxysome genes in other *Ca.* Halomarinus genomes (Fig. 2), none were annotated in the LSUCC0096 genome. Conversely, the LSUCC0096 *cbbQ* and *cbbO* genes had no matching orthologs in any of the other *Ca.* Halomarinus genomes. Thus, the LSUCC0096 RuBisCO is in a unique gene neighborhood and likely has a separate evolutionary history from the other *Ca.* Halomarinus RuBisCO genes.

Multiple pathways for phosphoglycolate salvage have recently been investigated in the model chemolithoautotrophic organism *Cupriavidas necator* H16 (77). We found annotated genes supporting the presence of the C2 cycle (*glyA*, *hprA*, and associated aminotransferases) and the malate cycle (*aceB*, *maeB*, pyruvate dehydrogenase *aceEF*) in *Ca.* Halomarinus genomes, but genes for the oxalyl-CoA decarboxylation route, as well as the *gcl* gene for the glycerate pathway, appear to be missing.

### N, P, and S

*Ca.* Halomarinus uses the PII nitrogen response system and 24 of 25 genomes had the *amtB* ammonia transporter (14 genomes had two copies), with ten genomes containing complete urea transporter genes (*urtABCDE*), and others with partial transporters (Fig. 2, Table S1). Urease alpha, beta, and gamma subunit genes were conserved in HIMB30 and LSUCC0096 and five other genomes across both subclades (Fig. 2, Table S1), with partial urease genes found in more genomes. Complete or partial phosphate transporter genes (*pstABC*) were conserved across 17 genomes, and ten in both subclades were predicted to transport phosphonate (*phnCDE*) as well (Fig. 2). However, the *phn* C-P lyase genes were present exclusively in a subset of seven subclade I genomes. Both subclades had predicted genes for sulfide oxidation, and the sulfite dehydrogenase had variable distribution across the subclades as well (Fig. 2). We also found *sox* genes for thiosulfate oxidation in both subclades, with the exception of the species cluster *Ca.* H. estuariensis in subclade I (Fig. 2), and all genomes contained at least one copy of a sulfite exporter (*tauE*) (Table S1). Thus, subclades I and II were predicted to carry out sulfide- and thiosulfate-based chemolithotrophy. DMSP demethylation and synthesis genes were missing from all genomes, although three genomes had predicted DMSP lyases (Fig. 2).

### Other features

The majority of *Ca.* Halomarinus genomes contained biosynthesis pathways for the bulk of essential amino acids, but none of the genomes contained genes for phenylalanine biosynthesis (Fig. 2). Thus, it appears this auxotrophy is conserved across the clade. Branched-chain and polar amino acid ABC transporters were present in the majority of genomes, as was a glycine betaine/proline ABC transporter (Table S1). B vitamin biosynthesis was limited. Thiamin (B1) and riboflavin (B2) biosynthesis pathways were partially complete in genomes from both subclades (Fig. 2). Most genomes had predicted *ribBAHE* genes for riboflavin synthesis from ribulose-5P. Fourteen genomes contained the *thiXYZ* transporter for hydroxymethylpyrimidine (HMP) and 24 and 23 contained *thiD* and *thiE*, respectively (Fig. 2, Table S1). Thus, *Ca.* Halomarinus may synthesize thiamin from imported HMP. *Ca.* Halomarinus genomes appeared auxotrophic for biotin (B7), but possessed the biotin transporter component *bioY*. These organisms additionally had only partial pathways for pantothenate (B5), pyridoxine (B6), and folate (B9). No genes were present for nicotinamide/nicotinate (B3) biosynthesis, although NAD+ biosynthesis was intact. LSUCC0096, the most complete genome, was the only genome with a predicted *btuB* transporter component for cobalamin (B12). Most also contained genes for transport of ferric iron (*afuABC* - 23/25), copper (*copA* - 14/25), tungstate (*tupABC* - 15/25), zinc (*znuABC* - 12/25), and chromate (*chrA* - 12/25) (Fig. 2, Table S1). A small subset of genomes, including HIMB30 and LSUCC0096, contained all the genes for molybdate (*modABC* - 4/25), and iron complex transport (*fhuDBC* - 2/25), although all but three genomes had a predicted *fhuC* (Fig. 2, Table S1). None of the genomes with ureases contained annotated *nik* transporters for nickel, despite it being the required cofactor for urease. Thus, nickel may be obtained by promiscuous activity from one of the other ABC transporters in the genome, or they may be mis-annotated (78).

Eighteen genomes from both subclades had genes for flagellar biosynthesis, and so we predict *Ca.* Halomarinus cells to be motile (Fig. 2). Consistent with a sodium-motive force in OM252 cells, many of genomes contained sodium symporters for phosphate (8/25), acetate (15/25), and melibiose (25/25), as well as sodium antiporters for calcium (*yrbG* - 23/25) and protons (*nhaA* - 15/25) (Fig. 2, Table S1). The latter may provide a useful means for converting the proton motive force generated by proteorhodopsin to a sodium motive force in some strains. There was also a proton-chloride antiporter (*clcA*) conserved in 23 genomes. We found peroxiredoxin in 21 genomes and catalase (*katG*) in six genomes spanning both subclades as well (Fig. 2, Table S1). Finally, almost all genomes had *phbBC* genes to synthesize (and degrade) poly-ß-hydroxybutyrate (PHB), and two associated phasin genes were found in many genomes as well.

### Morphology and growth characteristics of LSUCC0096

Cells of LSUCC0096 were curved-rod/spirillum-shaped, approximately 1.5 μm long and 0.2-0.3 μm wide (Figs. 3, S18). We also found evidence of a flagellum (Figs. 3 inset, S18BD), corroborating genomic predictions (above). Coastal Louisiana experiences dramatic shifts in salinity owing to a large number of estuaries in the region and tidal forcing through barrier islands, marshes, and different delta formations (79). Our previous 16S rRNA gene data suggested that OM252, and specifically the ASV5512 that matched LSUCC0096, had a euryhaline lifestyle, being found across a range of salinities but having greater prevalence in salinities above 12 (12). Therefore, we examined the salinity tolerance of LSUCC0096 through two complimentary methods: first, by altering only the concentration of NaCl in the medium, and second, by changing all major ion concentrations proportionally (Fig. 4A). LSUCC0096 grew in salinities between 5.79 to 63.5, with a maximum growth rate of 0.23 (+/− 0.01) doublings/hour at 11.6 under the proportional scheme. We detected no growth at salinities of 0.36 or 1.45 (Fig. S19A). Although there was an overlap of salinities from 8.66 to 34.8 between the two experiments, the growth rates were higher when the ion concentrations were altered proportionally compared to when only the concentration of NaCl was altered (Fig. 4A). LSUCC0096 also grew between 12°C and 35°C in the isolation medium JW1 with a maximum growth rate of 0.36 (+/− 0.06) doublings/hour at 35°C and a “typical” growth rate of 0.19 (+/− 0.03) at 25°C (Fig. 4B). We did not detect growth at 4°C or 40°C (Fig. S19B).

**Figure 3.**
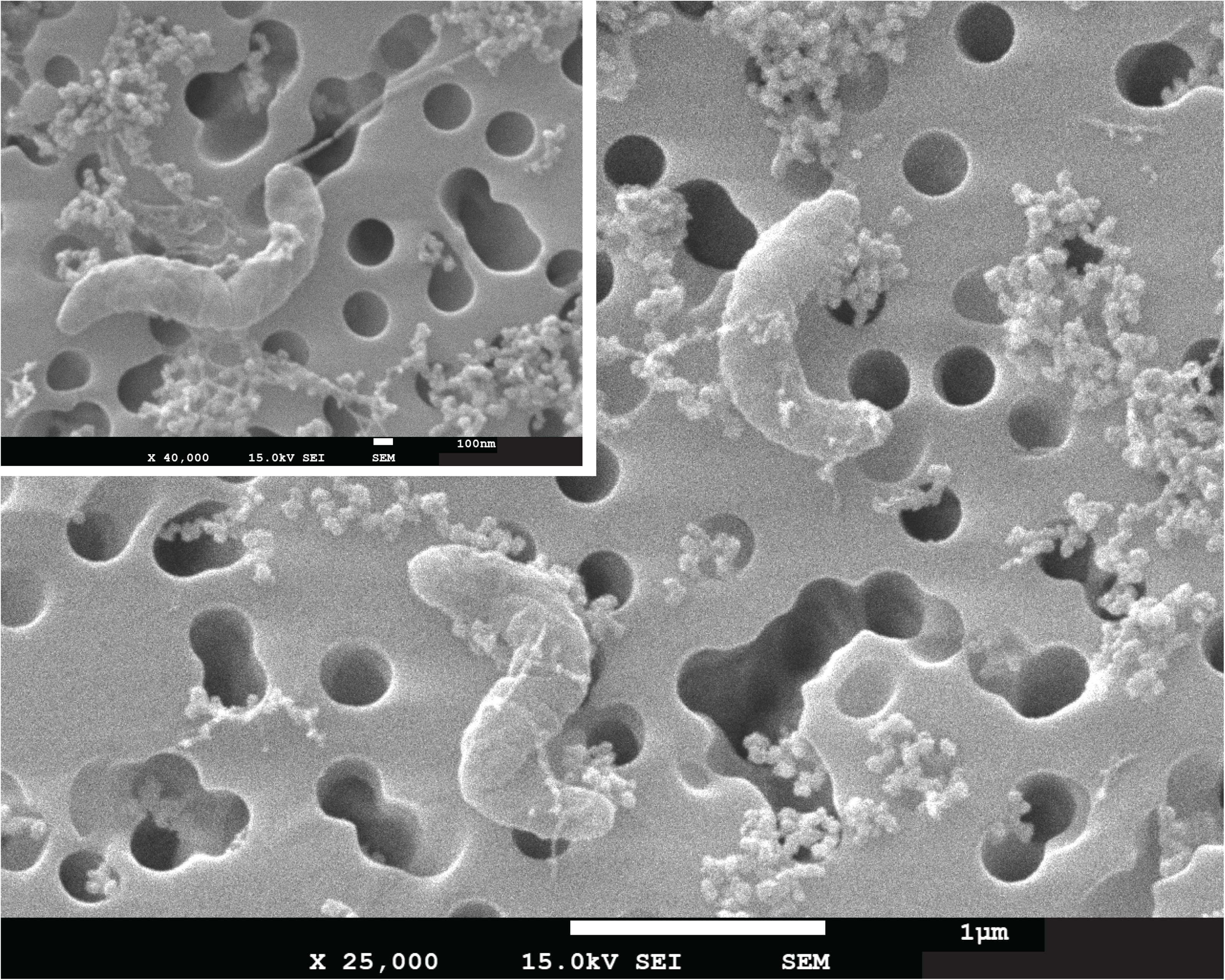
Scanning electron micrographs of LSUCC0096. Main-25,000x magnification of two cells on a 0.2 μm filter, scale bar = 1 μm. Inset-40,000x magnification of a dividing cell to focus on possible polar flagellum in the upper pole, scale bar = 100 nm.

**Figure 4.**
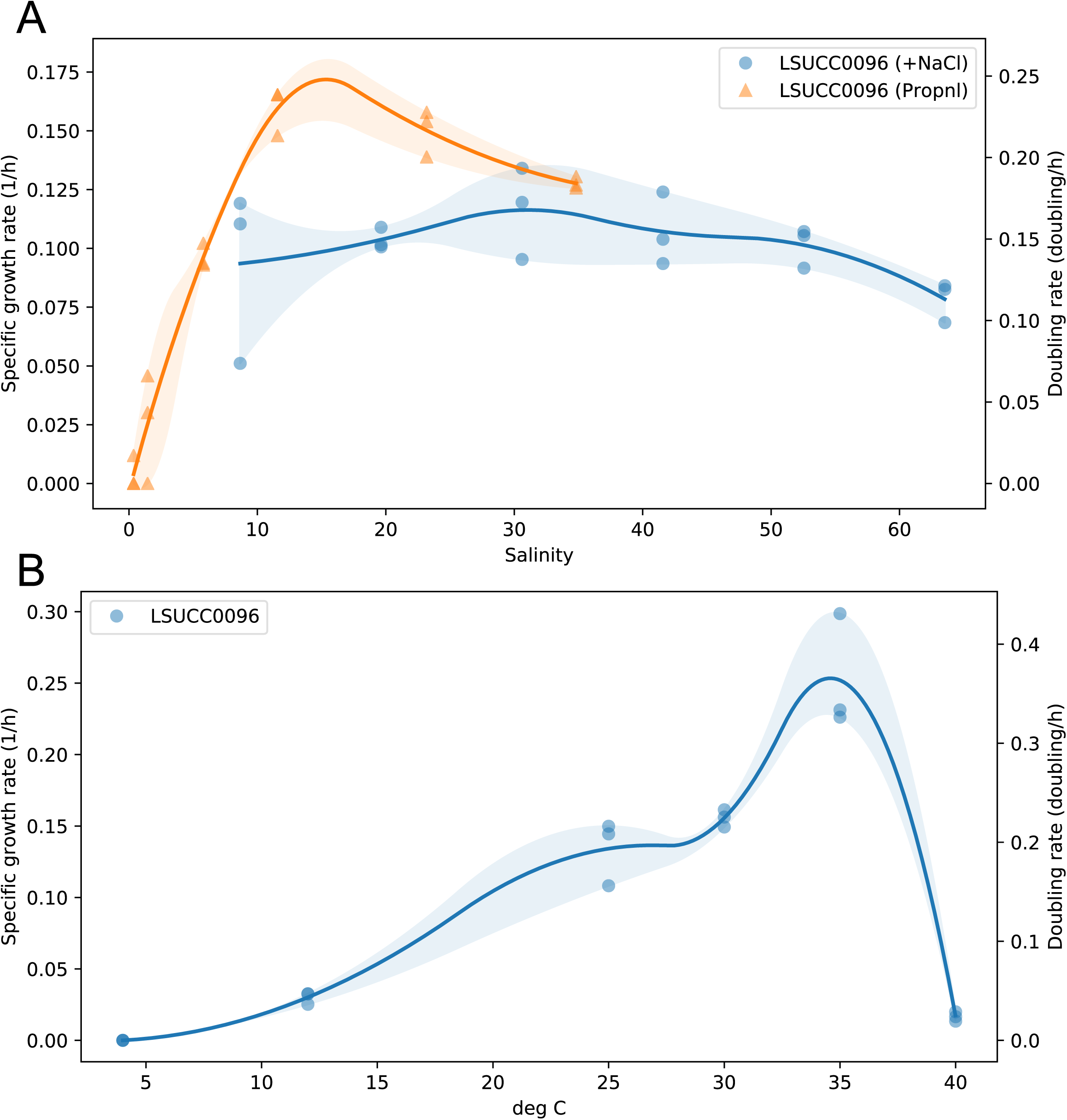
Salinity and temperature growth ranges for LSUCC0096. A) Specific growth rates and doubling times according to variable salinity based on proportional dilution of major ions (orange) or changing only NaCl concentration (blue) within the media. B) Specific growth rates and doubling times according to temperature.

### Thiosulfate-dependent chemolithoautotrophic growth

We tested the ability of LSUCC0096 to grow under chemolithoautotrophic conditions with thiosulfate as the sole electron donor. We measured growth of LSUCC0096 across four consecutive transfers in modified JW1 medium with no added organic carbon (other than trace quantities of vitamins) and 100 μM thiosulfate. Inorganic carbon was present as bicarbonate (10 mM), used as the medium buffer (10). Growth curves from the fourth growth cycle are presented in Figure 5. When grown under strict chemolithoautotrophic conditions, LSUCC0096 increased in cell density more than two orders of magnitude in a typical logarithmic growth pattern, albeit more slowly, and to a lower cell density, than when grown in chemoorganoheterotrophic conditions (Fig. 5). The positive controls from the fourth transfer had an average growth rate of 0.20 +/− 0.01 doublings/hour, which is similar to growth rates found under normal growth conditions (Fig. 4), whereas the experimental replicates had a much slower average growth rate of 0.07 +/− 0.01 doublings/hour (Fig. S19C). Growth yields under chemolithoautotrophic conditions were roughly 68% of that under chemoorganoheterotrophic conditions (1.26 +/− 0.85 × 10^6^ vs. 1.85 +/− 1.29 × 10^6^ cells/ml), although the variance overlapped. The overall Gibbs energies, Δ*G_r_*, of organic carbon and thiosulfate oxidation under the experimental conditions were −113.2 kJ (mol e^−^)^−1^ and −100.9 kJ (mol e^−^)^−1^, respectively. We observed limited growth in the negative controls, but these were inconsistent and at much lower rates than for the experimental conditions (Fig. S19C). It is possible that storage compounds like PHB were not fully exhausted after four successive transfers, thus supplying the necessary energy and carbon for limited additional growth (80). Nevertheless, in conjunction with the genomic data, our experimental results provide strong evidence that LSUCC0096 is capable of oxidizing thiosulfate as a facultative chemolithoautotroph.

**Figure 5.**
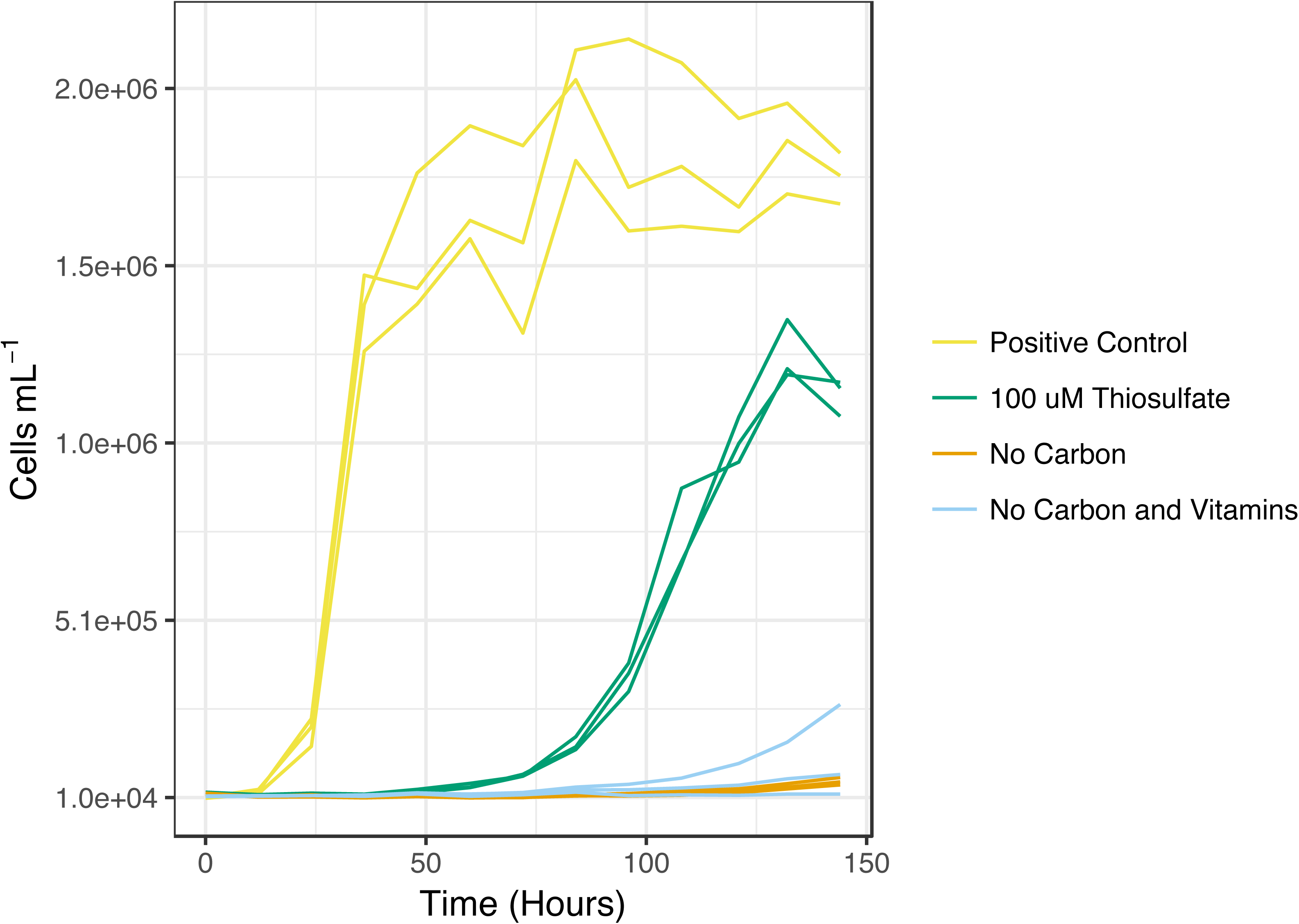
Thiosulfate-based chemolithoautotrophic growth in LSUCC0096. Cell numbers plotted against time for growth in chemolithoautotrophic conditions with thiosulfate as the sole electron donor (green), compared with typical heterotrophic medium (yellow), and no carbon (orange) and no carbon/vitamin (blue) controls. Curves depict growth after four consecutive transfers.

## Discussion

We comprehensively examined the distribution, genomic diversity, and taxonomy of OM252 bacterioplankton using 25 genomes from two pure cultures (including our recently isolated strain LSUCC0096), 7 SAGs, and 16 MAGs. These organisms were generally characterized by genomes in the 2.2 Mb range, with ~ 49% GC content and coding densities ~ 92%, although the two most complete genomes, from isolates HIMB30 and LSUCC0096, had 94 and 95% coding density, respectively. Thus, OM252 genomes are slightly larger and less streamlined than SAR11 genomes, but smaller than most Roseobacter spp. (81–83).

Images of strain LSUCC0096 indicate these cells are curved-rod/spirillum-shaped (Figs. 3, S18) and somewhat larger than typical SAR11 cells (84). LSUCC0096 cells were also narrower and longer than *Litoricola marina* and *Litoricola lipolytica*, which were described as short rods, with no mention of curvature (26, 73). It remains to be seen if LSUCC0096 morphology is conserved throughout the OM252 clade. *L. marina* and *L. lipolytica* were also reported to be non-motile (26, 73), whereas OM252 genomes contain flagellar genes and we found evidence of a polar flagellum in LSUCC0096 (Figs. 3 inset, S18BD).

Phylogenetics with 16S rRNA genes and concatenated single-copy marker genes, as well as ANI comparisons, corroborated the sister relationship of OM252 with the genus *Litoricola*, and defined two major subclades and several species boundaries within OM252. Currently, HIMB30 and several MAGs used in this study are classified as a *Litoricola* species in the Genome Taxonomy Database (GTDB). However, based on the depth of branching between *Litoricola* and OM252 within our trees, the ANI values among OM252 genomes, the pairwise 16S rRNA gene identities within the group and between OM252 and *Litoricola* (72), and the substantial difference in GC content between OM252 and *Litoricola* (47-51% vs. 58-60% (26, 73), respectively), we argue here for distinguishing OM252 as a separate genus, which we propose as *Candidatus* Halomarinus, gen. nov.. Our whole genome phylogeny is consistent with the current placement of *Ca.* Halomarinus within the Litoricolaceae, and for the Litoricolaceae within the Pseudomonadales, as currently defined in GTDB (https://gtdb.ecogenomic.org/, accessed February 2021). Poor branch support at the internal nodes grouping Litoricolaceae with SAR86, and that group within the remainder of the Pseudomonadales, precludes us from commenting on the likely position of Litoricolaceae in that Order (Figs. 1, S3).

*Ca.* Halomarinus bacteria are globally distributed in marine surface waters, and some strains can be found in bathy- and abyssopelagic depths. The *Ca.* H. estuarensis species cluster was predominant in the San Francisco Bay estuary (Fig. S14), but recruited poorly from open ocean samples (Fig. S5). Thus, certain species may have more restricted biogeography than others. Increases in taxon selection and additional coastal and estuarine metagenomic sampling will improve these types of assessments. We demonstrated that strain LSUCC0096 can grow over a wide range of salinities (5.77-63.6), although it appears to be adapted for brackish conditions with an optimal growth salinity of 11.6 (Fig. 4A). Our coastal 16S rRNA gene data previously demonstrated that the ASV matching LSUCC0096 had maximum abundances in salinities between 12-21 (12) and recruitment data from the San Francisco Bay Estuary system (Fig. S12) supports the hypothesis that LSUCC0096 represents a brackish water specialist, but is incapable of growth in fresh water (Fig. S18). Nevertheless, this organism also recruited reads from all over the globe (Fig. S4) and displayed considerable halotolerance. Although we measured growth at up to 5% NaCl (calculated salinity 63.5), we did not actually find the maximum salinity beyond which the cells could not grow (Fig. 4A). The extensive salinity tolerance of strain LSUCC0096 corroborates culture-independent detection of the OM252 clade in very salty environments like the Salton Sea (18) and Spanish salterns (19), as well as their cosmopolitan distribution in the global oceans (Fig. S4). Future work on additional *Ca.* Halomarinus isolates will expand our understanding of the halotolerance and optimal salinities for the various other species in the group. Thermal tolerances were more pedestrian, with LSUCC0096 exhibiting a mesophilic temperature growth range.

Comparative genomics predicted that *Ca.* Halomarinus spp. are obligate aerobes with the capacity for both chemoorganoheterotrophic and sulfur-oxidizing chemolithotrophic metabolism. In support of these predictions, both existing isolates, strains HIMB30 and LSUCC0096, were isolated under aerobic, chemoorganoheterotrophic growth conditions (10, 22). We predict that *Ca.* Halomarinus spp. can utilize TCA cycle intermediates, some sugars, and possibly amino acids as carbon and energy sources, although direct characterization of the suite of compounds that can be used needs further investigation. Given the possibility for utilization of the storage compound PHB, either as both an energy and carbon source or as an energy source in conjunction with RuBisCO-based carbon fixation (80), future experiments would require a CO_2_-free headspace, an alternative buffer to the bicarbonate used in JW1, and probably five or more successive growth cycles to eliminate storage compounds.

*Ca.* Halomarinus appears to subsist on ammonia or urea as nitrogen sources, with phosphate as the primary source of phosphorous, and phosphonates as possible substitutes for some strains, similarly to SAR11 (81, 85). However, many Pelagibacterales lack the PII system (86) that is present in OM252. *Ca.* Halomarinus spp. can likely synthesize most amino acids except phenylalanine. We predicted B vitamin synthesis is limited to riboflavin and thiamin via *thiDE* after import of HMP. Thiamin biosynthesis after HMP import is also similar to the Pelagibacterales (87), with the major difference being the presence of the *thiXYZ* HMP transporter in OM252. We also predict *Ca.* Halomarinus spp. utilize ferric iron and may additionally interact with copper, tungstate, zinc, and chromate. Thus, they have a similarly restricted set of metal transporters as SAR11, and far fewer than many Roseobacter spp. (88).

Most *Ca.* Halomarinus genomes had predicted genes for oxidation of reduced sulfur compounds, and subclade I organisms additionally had genes for autotrophy via the CBB cycle and RuBisCO. Corroborating these predictions, we demonstrated that the subclade I representative LSUCC0096 could grow for successive transfers under strict chemolithoautotrophic conditions with thiosulfate as the sole electron donor and bicarbonate as the sole available carbon source (excepting vitamins). The ability to switch between autotrophic and heterotrophic metabolism also has biogeochemical relevance because it means that these organisms can switch between serving as inorganic carbon sources and sinks. This behavior has implications for modeling marine carbon cycling since these organisms cannot be simply classified as heterotrophic. The pervasiveness of facultative lithoautotrophy among putative “heterotrophic” lineages deserves further investigation.

The relevance of facultative lithoautotrophy to both the carbon and sulfur cycles also places emphasis on understanding what may control these different lifestyles in nature. The experimental data from strain LSUCC0096 (growth rates and lag times after repeated transfers) suggest heterotrophic growth will always be favored to lithoautotrophic growth in *Ca*. Halomarinus subclade I strains, and calculations indicate that this probably arises due to the energetics of anabolism rather than catabolism. The energy available from organic carbon oxidation was only about 12% greater than thiosulfate oxidation, yet the growth rate was nearly three times greater in the heterotrophic experiment. This divergence could be explained by the much larger difference in the energetics of biomolecule synthesis when the starting materials are inorganic compounds such as CO_2_ and NH_4_^+^ versus a suite of organic compounds. The energetics of protein synthesis provides an illustrative example since bacterial cells are approximately 50% protein (89). If an environment is replete with amino acids, then microorganisms need only to obtain and polymerize them to build proteins. The Gibbs energy of peptide bond formation is ~40 kJ per mol^−1^ (90). However, if the organisms must first synthesize amino acids *de novo* before polymerization, the cost is much greater. For instance, the Δ*G_r_* of alanine synthesis from CO_2_ and NH_4_^+^ in an oxidizing environment is 1,380 kJ per mol^−1^, and for more complex amino acids such tryptophan, it is 5,320 kJ per mol^−1^ (91). Therefore, the LSUCC0096 cells in the thiosulfate experiment had to dedicate a larger *flux* of their catabolic energy to biomolecule synthesis than the heterotrophs who were essentially given all of the starting materials. These conclusions suggest that thiosulfate-based chemolithoautotrophy is utilized in nature when organic compound concentrations become limiting. However, more research is required to understand whether there is a complete metabolic switch to autotrophy, and if so, whether it is strictly controlled by the relative availability of growth substrates or if some additional regulation is involved.

Another intriguing mystery is the source and temporal availability of thiosulfate and other reduced sulfur compounds in natural marine systems. Abiotic oxidation of sulfide results in the generation of stable thiosulfate in seawater, an effect that was enhanced by the presence of trace metals like Fe, Pb, and Cu (92). Thus, any source of sulfide could theoretically lead to production of thiosulfate if the sulfide is not first consumed by other microbes. Alternatively, thiosulfate may occur as a transient intermediate as a direct result of microbial metabolism, a process labeled the “thiosulfate shunt” in sedimentary systems (93). If it could escape into the oxic water column from systems with low or no oxygen, organisms like *Ca.* Halomarinus may be able to harvest energy from thiosulfate originating from cryptic sulfur cycling processes near OMZs and sinking particles (94–96), or possibly also shallow sediments, where thiosulfate can persist in micromolar concentrations (97, 98). Researchers have been isolating thiosulfate-utilizing bacteria from seawater for over a century (99), and it has long been known that thiosulfate-oxidizers occupy the marine water column and can connect this metabolism to carbon-fixation activity (100–102). More recently, we have also learned that *sox* genes are common in the oxic marine water column (94, 103, 104). Indeed, many marine prokaryotes contain these genes and thiosulfate oxidation can be used to stimulate growth under mixotrophic conditions and even anapleurotic carbon-fixation (see, for example, (105–108)). Thus, a variety of microorganisms from oxic marine waters are poised for thiosulfate-based chemolithotrophy and sometimes autotrophy. The circumstances and controls on reduced inorganic sulfur compound use by obligate aerobes like *Ca.* Halomarinuin the oxic water column requires further study.

### Description of *Halomarinus*, gen. nov.

*Halomarinus* (Ha.lo.ma.ri.nus G. masc. n. *halo* salt, sea; L. masc. adj. *marinus*, of the sea; N.L. masc. n. *Halomarinus* salty, seagoing, in reference to the marine habitat and high salinity tolerance of the organisms).

Aerobic, chemoorganoheterotrophic and chemolithotrophic, with sodium-translocating NADH dehydrogenases, capable of glycolysis, gluconeogenesis, and possessing a complete TCA cycle. Has genes for motility via flagella. Possesses the PII-dependent nitrogen response system and genes for ammonia, phosphate, ferric iron, tungstate, copper, zinc, chromate transport. Has genes for synthesizing histidine, arginine, lysine, serine, threonine, glutamine, cysteine glycine, proline, methionine, isoleucine, leucine, tryptophan, tyrosine aspartate, glutamate, but are phenylalanine auxotrophs. Genes for synthesis of riboflavin (vitamin B2) and thiamine (vitamin B1) from HMP. Auxotrophic for vitamins B3, B5, B6, B7, B9, B12. Has genes for poly-ß-hydroxybutyrate production and degradation and peroxiredoxin. Estimated complete genome sizes between 1.49 - 2.68 Mbp, GC content between 47 - 51%, coding densities between 82-96%.

The type species is *Candidatus* Halomarinus kaneohensis.

### Description of *Candidatus* Halomarinus kaneohensis, sp. nov.

*Candidatus* Halomarinus kaneohensis (ka.ne.o.hen.sis N.L. n. *Kāne*ʻ*ohe,* a bay on the island of Oahu, HI, USA, from where the strain was isolated).

In addition to the characteristics for the genus, it has the following features. Has proteorhodopsin and retinal biosynthesis genes. Has a predicted *cbb_3_*-type cytochrome c oxidase, genes for the glyoxylate shunt, urease, and the *coxMSL* aerobic carbon monoxide dehydrogenase genes. Is predicted to be capable of thiosulfate and sulfide oxidation, as well as autotrophy via the Calvin-Benson-Bassham cycle.

The type strain, HIMB30^T^, was isolated from seawater collected in Kāneʻohe Bay, Oahu, HI, U.S.A. (21.460467, −157.787657) (22). The genome sequence for HIMB30^T^ is available under NCBI BioProject <u>PRJNA47035</u>. Estimated complete genome size of 2.26 Mbp, GC content of 50% from genome sequencing. The culture is maintained in cryostocks at the University of Hawaiʻi at Mānoa by M.S. Rappé. We provide the *Candidatus* designation since the culture has not been deposited in two international culture collections and therefore does not satisfy the naming conventions of the International Code of Nomenclature for Prokaryotes (ICNP) (109). However, the characterization here is more than sufficient for naming recognition via genomic type material (110, 111).

### Description of *Candidatus* Halomarinus pommedorensis, sp. nov.

*Candidatus* Halomarinus pommedorensis (pomme.d.or.en.sis N.L. n. *Pomme d’Or*, a bay in southern Louisiana, U.S.A., from where the strain was isolated).

In addition to the characteristics for the genus, it has the following features. Cells are curved-rod/spirillum-shaped, ~ 1.5 μm × 0.2-0.3 μm. Halotolerant, being capable of growth in salinities between 5.8 and at least 63.4, but not at 1.5 or below. Mesophilic, being capable of growth at temperatures between 12 and 35°C but not at 4 or 40°C. Has a maximum growth rate at 35°C in the isolation medium JW1 of 0.36 (+/− 0.06) doublings/hour. Has a predicted *cbb_3_*-type cytochrome c oxidase and genes for the glyoxylate shunt, urease, and cobalamin transport. Has predicted genes for thiosulfate and sulfide oxidation, as well as autotrophy via the Calvin-Benson-Bassham cycle. Grows under thiosulfate-oxidizing chemolithoautotrophic conditions at 0.07 (+/− 0.01) doublings/hour.

The type strain, LSUCC0096^T^, was isolated from seawater collected at Bay Pomme d’Or, Buras, LA, U.S.A. (29.348784, −89.538171) (10). The GenBank accession number for the 16S rRNA gene of LSUCC0096^T^ is <u>KU382366.1</u>. The genome sequence is available under BioProject <u>PRJNA551315</u>. The culture is maintained in cryostocks at the University of Southern California by J.C. Thrash. Estimated complete genome size of 2.01 Mbp, GC content of 49% from genome sequencing. We provide the *Candidatus* designation since the culture has not been deposited in two international culture collections and therefore does not satisfy the naming conventions of the ICNP (109). However, the characterization here is more than sufficient for naming recognition via genomic type material (110, 111).

### Description of *Candidatus* Halomarinus estuariensis, sp. nov.

*Candidatus* Halomarinus estuariensis (es.tu.ar.i.en.sis L. masc. adj. *estuarine*, based on its relative abundance in an estuary).

In addition to the characteristics for the genus, it has the following features. Has proteorhodopsin and retinal biosynthesis genes. Has an additional *cbb_3_*-type cytochrome c oxidase. Is predicted to be capable of sulfide oxidation, as well as autotrophy via the Calvin-Benson-Bassham cycle, but not thiosulfate oxidation. Has predicted genes for the glyoxylate shunt, as well as D-galacturonate epimerase. Some strains have C-P lyase and DMSP lyase genes. Estimated complete genome sizes between 1.98 and 2.68 Mbp, GC content between 47 - 49% from genome sequencing.

We provide the Candidatus designation since this species has not yet been cultivated. Genomes were reconstructed from metagenomic sequencing.

## Data Availability

The LSUCC0096 genome is available on NCBI under BioProject number <u>PRJNA551315</u>, and IMG under Taxon ID 2639762503. The raw reads from which LSUCC0096 was assembled are available at the NCBI SRA under accession number <u>SRR9598636</u>. Metagenomic sequences from the San Francisco Bay estuary were used with permission from Dr. Christopher A. Francis and are available on NCBI under the following SRA accessions: SRR7130817, SRR7130819, SRR7130820, SRR7130903, SRR7131305, SRR7131306, SRR7132116, SRR7132117. Cryostocks and/or live cultures of strains LSUCC0096 and HIMB30 are available upon request.

## Acknowledgements

This work was supported by a Louisiana Board of Regents grant [LEQSF(2014-17)-RD-A-06], a Simons Early Career Investigator in Marine Microbial Ecology and Evolution Award, and NSF Biological Oceanography Program grants (OCE-1747681 and OCE-1945279) to JCT. The authors also thank Dr. Christopher A. Francis for use of the San Francisco Bay metagenomic dataset.

