## Supplemental Information for "Ecophysiology of the cosmopolitan OM252 bacterioplankton (Gammaproteobacteria)"

**Supplemental Figures**

Figure S1. 16S rRNA gene maximum-likelihood phylogeny of the Gammaproteobacteria with near neighbors from the NCBI RefSeq database. OM252 clade is highlighted in grey. Bootstrap support values (n=1000) are indicated at the nodes. Scale bar = changes per position.

Figure S2. 16S rRNA gene maximum-likelihood phylogeny of the Gammaproteobacteria with near neighbors from the NCBI nt database, including clone library sequences. OM252 clade is highlighted in grey. Bootstrap support values (n=1000) are indicated at the nodes. Scale bar = changes per position.

Figure S3. Expanded phylogenomic tree of the OM252 clade (same as Figure 1 but with all branches shown). Maximum-likelihood tree based on 78 concatenated single-copy genes within the Pseudomonadales (as designated by GTDB) and selected other Gammaproteobacteria. Final alignment = 29,631 amino acid positions. Families designated by GTDB within the Pseudomondales are indicated, with shading for the OM252 clade. Species designated in this study are highlighted in red and orange text. Values at nodes indicate bootstrap support (n=1000), scale indicated changes per position.

Figure S4. Global visualization of metagenomic recruitment to the OM252 genomes from the Tara Oceans, Biogeotraces, and Malaspina datasets. Genome name is indicated in the boxes above each map. $Log_{10}$-transformed RPKM values are indicated via the bubble size in the key.

Figure S5. Distribution of all metagenomic recruitment to the OM252 genomes from the Tara Oceans, Biogeotraces, and Malaspina datasets. Box plots depict the distribution of the $Log_{10}$-transformed RPKM values for every site (n=605) per genome. Outlying points are plotted outside the boxplot range.

Figure S6. Diversity Principal Coordinates Analysis (PCoA) from unifrac distances accounting for recruited OM252 reads from 588 metagenomic samples. Distances are weighted relative to the phylogenetic relationship of each OM252 community representative. Scatters are individual community abundances in a given metagenomic sample. PC1 is a primary indication of dissimilarity and is measured horizontally. PC2 is a second indication of dissimilarity and is measured by transparent depth. PC3 is a tertiary indication of dissimilarity and is measured vertically. ANOSIM correlation statistics are shown for each PCoA. All ANOSIM p-tests passed with a standard value (p-val=0.001). The skbio.diversity algorithm was retrofitted for specific use of the OM252 dataset using (https://github.com/thrash-lab/diversity_metrics).

Figure S7. Per-site recruitment vs. salinity. $Log_{10}$-transformed RPKM values are plotted against salinity for every site in the Tara Oceans, Biogeotraces, and Malaspina datasets containing that data. Genome name is indicated in the boxes above each map. Linear regressions and 95% confidence intervals are plotted in grey.

Figure S8. Per-site recruitment vs. temperature. $Log_{10}$-transformed RPKM values are plotted against temperature for every site in the Tara Oceans, Biogeotraces, and Malaspina datasets containing that data. Genome name is indicated in the boxes above each map. Linear regressions and 95% confidence intervals are plotted in grey.

Figure S9. Per-site recruitment vs. depth. $Log_{10}$-transformed RPKM values are plotted against depth for every site in the Tara Oceans, Biogeotraces, and Malaspina datasets. Genome name is indicated in the boxes above each map. Non-linear regressions and 95% confidence intervals are plotted in grey.

Figure S10. Distribution of sampled sites, by latitude, within the Tara Oceans, Biogeotraces, and Malaspina datasets. A) Histogram of sampling sites by latitude, with individual datasets colored according to the key. B) Histogram of sampling sites by latitude, with density of sites overlaid in pink.

Figure S11. Per-site recruitment vs. latitude. $Log_{10}$-transformed RPKM values are plotted against latitude for every site in the Tara Oceans, Biogeotraces, and Malaspina datasets. Genome name is indicated in the boxes above each map. Non-linear regressions and 95% confidence intervals are plotted in grey.

Figure S12. Local visualization of metagenomic recruitment to the OM252 genomes from the northern Gulf of Mexico, San Pedro Basin, and San Francisco Bay Estuary datasets. Genome name is indicated in the boxes above each map. $Log_{10}$-transformed RPKM values are indicated via the bubble size in the key.

Figure S13. Distribution of all metagenomic recruitment to the OM252 genomes from the Gulf of Mexico, San Pedro Basin, and San Francisco Bay Estuary datasets. Box plots depict the distribution of the $Log_{10}$-transformed RPKM values for every site (n=26) per genome. Outlying points are plotted outside the boxplot range.

Figure S14. Distribution of metagenomic recruitment to the OM252 genomes from San Francisco Bay Estuary dataset specifically. Box plots depict the distribution of the $Log_{10}$-transformed RPKM values for every site (n=8) per genome. Outlying points are plotted outside the boxplot range.

Figure S15. Coastal recruitment vs. salinity. $Log_{10}$-transformed RPKM values are plotted against salinity for every site in the Gulf of Mexico, San Pedro Basin, and San Francisco Bay Estuary datasets containing that data. Genome name is indicated in the boxes above each map. Linear regressions and 95% confidence intervals are plotted in grey.

Figure S16. Complete metabolic reconstruction of the OM252 clade. Same as Figure 2, but including genes and pathways absent in all genomes. Heatmap displays gene and pathway content according to the scale on the right. Subgroups of processes and key metabolic pathways are highlighted for ease of viewing.

Figure S17. Maximum-likelihood phylogeny of the RuBisCO large subunit gene. OM252 clade members are highlighted in green. Bootstrap support values (n=1000) are indicated at the nodes. Scale bar = changes per position.

Figure S18. Scanning electron micrographs of LSUCC0096. A and C- 25,000x magnification of cells on a 0.2 µm filter, scale bar = 1 µm. B and D- 40,000x magnification of individual cells to focus on possible polar flagella, scale bar = 100 nm. C and D are depicted in Figure 3 and reproduced here for ease of comparison with other images.

Figure S19. Growth data for LSUCC0096. Each box depicts datapoints of cells/mL vs. time in hours under different experimental conditions. Replicates are indicated in the inset boxes. Colored solid and dashed lines indicate linear regressions assigned by the sparse-growth-curve script to positive growth and death, respectively. A) Cell numbers vs. time (hours) by salinity, altered via dilution of the major ions ("Propnl") or by changing the NaCl concentration ("+NaCl"). B) Cell numbers vs. time (hours) by temperature. C) Cell numbers vs. time (hours) for the fourth consecutive transfer of the thiosulfate growth experiment. 100 uM ThioS- experimental treatment with 100 µM thiosulfate as the sole electron donor and bicarbonate as the carbon source; No C Neg- base medium with no carbon added; No C/Vit Neg- base medium with no carbon or vitamins added; Pos Contl- normal JW1 medium with all carbon sources and vitamins.

Figure S1

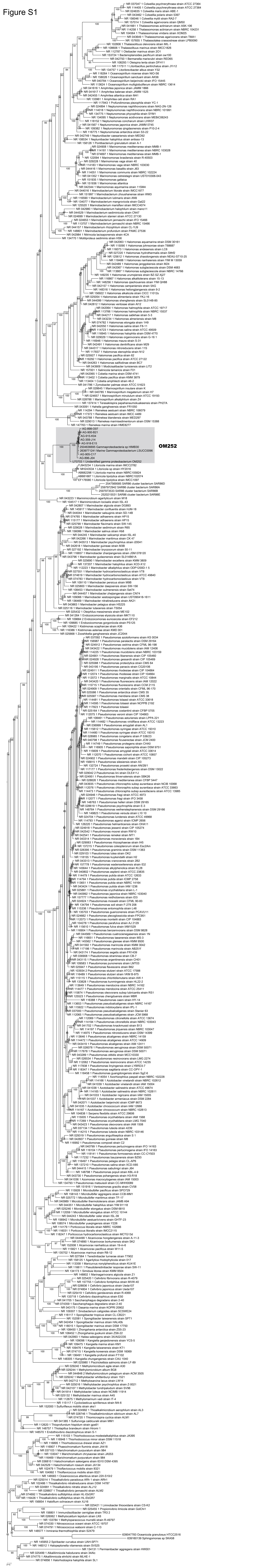

Figure S2

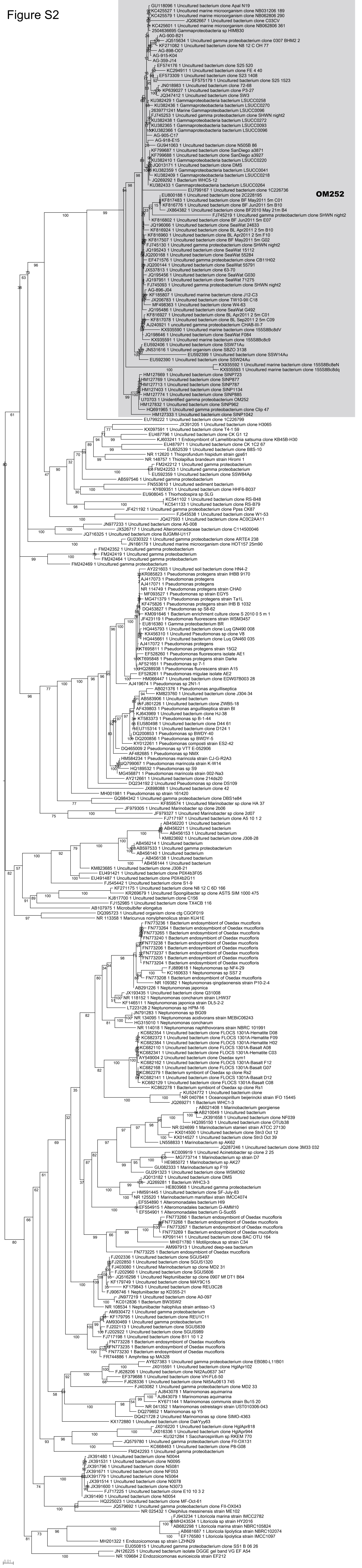

Figure S3

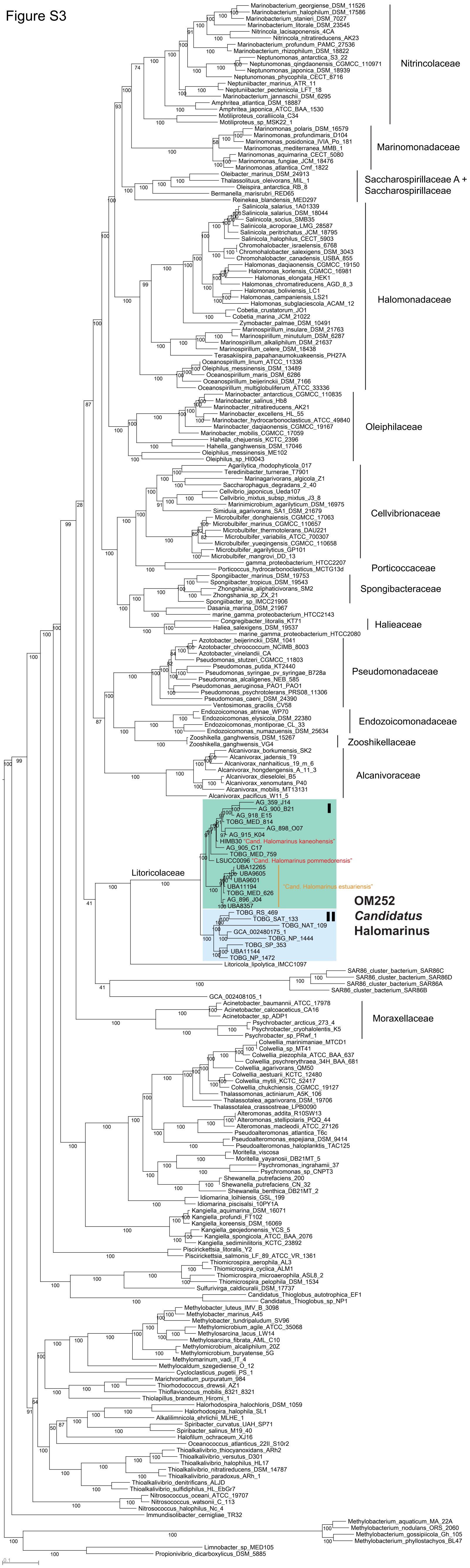

Marinobacterium_georgiense_DSM_11526
Marinobacterium_halophilum_DSM_17586
Marinobacterium_stanieri_DSM_7027
Marinobacterium_litorale_DSM_23545
Nitrincola_lacisaponensis_4CA
Nitrincola_nitratireducens_AK23
Marinobacterium_profundum_PAMC_27536
Marinobacterium_rhizophilum_DSM_18822
Neptunomonas_antarctica_S3_22
Neptunomonas_qingdaonensis_CGMCC_110971
Neptunomonas_japonica_DSM_18939
Neptunomonas_phycophila_CECT_8716
Neptuniibacter_marinus_ATR_11
Neptuniibacter_pectenicola_LFT_18
Marinobacterium_jannaschii_DSM_6295
Amphritea_atlantica_DSM_18887
Amphritea_japonica_ATCC_BAA_1530
Motiliproteus_coralliicola_C34
Motiliproteus_sp_MSK22_1
Nitrincolaceae
Marinomonas_polaris_DSM_16579
Marinomonas_profundimaris_D104
Marinomonas_posidonica_IVIA_Po_181
Marinomonas_mediterranea_MMB_1
Marinomonas_aquimarina_CECT_5080
Marinomonas_fungiae_JCM_18476
Marinomonas_atlantica_Cmf_1822
Marinomonadaceae
Oleibacter_marinus_DSM_24913
Thalassolituus_oleivorans_MIL_1
Oleispira_antarctica_RB_8
Bermanella_marisrubri_RED65
Reinekea_blandensis_MED297
Saccharospirillaceae A +
Saccharospirillaceae
Salinicola_salarius_1A01339
Salinicola_salarius_DSM_18044
Salinicola_socius_SMB35
Salinicola_acroporae_LMG_28587
Salinicola_peritrichatus_JCM_18795
Salinicola_halophilus_CECT_5903
Chromohalobacter_israelensis_6768
Chromohalobacter_salexigens_DSM_3043
Chromohalobacter_canadensis_USBA_855
Halomonas_daqiaonensis_CGMCC_19150
Halomonas_korlensis_CGMCC_16981
Halomonas_elongata_HEK1
Halomonas_chromatireducens_AGD_8_3
Halomonas_boliviensis_LC1
Halomonas_campaniensis_LS21
Halomonas_subglaciescola_ACAM_12
Cobetia_crustatorum_JO1
Cobetia_marina_JCM_21022
Zymobacter_palmae_DSM_10491
Marinospirillum_insulare_DSM_21763
Marinospirillum_minutulum_DSM_6287
Marinospirillum_alkaliphilum_DSM_21637
Marinospirillum_celere_DSM_18438
Terasakiispira_papahanaumokuakeensis_PH27A
Oceanospirillum_linum_ATCC_11336
Oleiphilus_messinensis_DSM_13489
Oceanospirillum_maris_DSM_6286
Oceanospirillum_beijerinckii_DSM_7166
Oceanospirillum_multiglobuliferum_ATCC_33336
Halomonadaceae
Marinobacter_antarcticus_CGMCC_110835
Marinobacter_salinus_Hb8
Marinobacter_nitratireducens_AK21
Marinobacter_excellens_HL_55
Marinobacter_hydrocarbonoclasticus_ATCC_49840
Marinobacter_daqiaonensis_CGMCC_19167
Marinobacter_mobilis_CGMCC_17059
Hahella_chejuensis_KCTC_2396
Hahella_ganghwensis_DSM_17046
Oleiphilus_messinensis_ME102
Oleiphilus_sp_HI0043
Oleiphilaceae
Agarilytica_rhodophyticola_017
Teredinibacter_turnerae_T7901
Marinagarivorans_algicola_Z1
Saccharophagus_degradans_2_40
Cellvibrio_japonicus_Ueda107
Cellvibrio_mixtus_subsp_mixtus_J3_8
Marinimicrobium_agarilyticum_DSM_16975
Simiduia_agarivorans_SA1_DSM_21679
Microbulbifer_donghaiensis_CGMCC_17063
Microbulbifer_marinus_CGMCC_110657
Microbulbifer_thermotolerans_DAU221
Microbulbifer_variabilis_ATCC_700307
Microbulbifer_yueqingensis_CGMCC_110658
Microbulbifer_agarilyticus_GP101
Microbulbifer_mangrovi_DD_13
Cellvibrionaceae
gamma_proteobacterium_HTCC2207
Porticoccus_hydrocarbonoclasticus_MCTG13d
Porticoccaceae
Spongiibacter_marinus_DSM_19753
Spongiibacter_tropicus_DSM_19543
Zhongshania_aliphaticivorans_SM2
Zhongshania_sp_ZX_21
Spongiibacter_sp_IMCC21906
Dasania_marina_DSM_21967
marine_gamma_proteobacterium_HTCC2143
Spongibacteraceae
Congregibacter_litoralis_KT71
Haliea_salexigens_DSM_19537
marine_gamma_proteobacterium_HTCC2080
Halieaceae
Azotobacter_beijerinckii_DSM_1041
Azotobacter_chroococcum_NCIMB_8003
Azotobacter_vinelandii_CA
Pseudomonas_stutzeri_CGMCC_11803
Pseudomonas_putida_KT2440
Pseudomonas_syringae_pv_syringae_B728a
Pseudomonas_alcaligenes_NEB_585
Pseudomonas_aeruginosa_PAO1_PAO1
Pseudomonas_psychrotolerans_PRS08_11306
Pseudomonas_caeni_DSM_24390
Ventosimonas_gracilis_CV58
Pseudomonadaceae
Endozoicomonas_atrinae_WP70
Endozoicomonas_elysicola_DSM_22380
Endozoicomonas_montiporae_CL_33
Endozoicomonas_numazuensis_DSM_25634
Endozoicomonadaceae
Zooshikella_ganghwensis_DSM_15267
Zooshikella_ganghwensis_VG4
Zooshikellaceae
Alcanivorax_borkumensis_SK2
Alcanivorax_jadensis_T9
Alcanivorax_nanhaiticus_19_m_6
Alcanivorax_hongdengensis_A_11_3
Alcanivorax_dieselolei_B5
Alcanivorax_xenomutans_P40
Alcanivorax_mobilis_MT13131
Alcanivorax_pacificus_W11_5
Alcanivoraceae
I
AG_359_J14
AG_900_B21
AG_918_E15
TOBG_MED_814
AG_898_O07
AG_915_K04
HIMB30 "Cand. Halomarinus kaneohensis"
AG_905_C17
TOBG_MED_759
LSUCC0096 "Cand. Halomarinus pommedorensis"
UBA12265
UBA9605
UBA9601
UBA11194
TOBG_MED_626
AG_896_J04
UBA8357
"Cand. Halomarinus estuariensis"
Litoricolaceae
II
TOBG_RS_469
TOBG_SAT_133
TOBG_NAT_109
GCA_002480175_1
TOBG_NP_1444
TOBG_SP_353
UBA11144
TOBG_NP_1472
Litoricola_lipolytica_IMCC1097
OM252
Candidatus
Halomarinus
SAR86_cluster_bacterium_SAR86C
SAR86_cluster_bacterium_SAR86D
SAR86_cluster_bacterium_SAR86A
SAR86_cluster_bacterium_SAR86B
GCA_002408105_1
Acinetobacter_baumannii_ATCC_17978
Acinetobacter_calcoaceticus_CA16
Acinetobacter_sp_ADP1
Psychrobacter_arcticus_273_4
Psychrobacter_cryohalolentis_K5
Psychrobacter_sp_PRwf_1
Moraxellaceae
Colwellia_marinimaniae_MTCD1
Colwellia_sp_MT41
Colwellia_piezophila_ATCC_BAA_637
Colwellia_psychrerythraea_34H_BAA_681
Colwellia_agarivorans_QM50
Colwellia_aestuarii_KCTC_12480
Colwellia_mytili_KCTC_52417
Colwellia_chukchiensis_CGMCC_19127
Thalassomonas_actiniarum_A5K_106
Thalassotalea_agarivorans_DSM_19706
Thalassotalea_crassostreae_LPB0090
Alteromonas_addita_R10SW13
Alteromonas_stellipolaris_PQQ_44
Alteromonas_macleodii_ATCC_27126
Pseudoalteromonas_atlantica_T6c
Pseudoalteromonas_espejiana_DSM_9414
Pseudoalteromonas_haloplanktis_TAC125
Moritella_viscosa
Moritella_yayanosii_DB21MT_5
Psychromonas_ingrahamii_37
Psychromonas_sp_CNPT3
Shewanella_putrefaciens_200
Shewanella_putrefaciens_CN_32
Shewanella_benthica_DB21MT_2
Idiomarina_loihiensis_GSL_199
Idiomarina_piscisalsi_10PY1A
Kangiella_aquimarina_DSM_16071
Kangiella_profundi_FT102
Kangiella_koreensis_DSM_16069
Kangiella_geojedonensis_YCS_5
Kangiella_spongicola_ATCC_BAA_2076
Kangiella_sediminilitoris_KCTC_23892
Piscirickettsia_litoralis_Y2
Piscirickettsia_salmonis_LF_89_ATCC_VR_1361
Thiomicrospira_aerophila_AL3
Thiomicrospira_cyclica_ALM1
Thiomicrospira_microaerophila_ASL8_2
Thiomicrospira_pelophila_DSM_1534
Sulfurivirga_caldicuralii_DSM_17737
Candidatus_Thioglobus_autotrophica_EF1
Candidatus_Thioglobus_sp_NP1
Methylobacter_luteus_IMV_B_3098
Methylobacter_marinus_A45
Methylobacter_tundripaludum_SV96
Methylomicrobium_agile_ATCC_35068
Methylosarcina_lacus_LW14
Methylosarcina_fibrata_AML_C10
Methylomicrobium_alcaliphilum_20Z
Methylomicrobium_buryatense_5G
Methylomarinum_vadi_IT_4
Methylocaldum_szegediense_O_12
Cycloclasticus_pugetii_PS_1
Marichromatium_purpuratum_984
Thiorhodococcus_drewsii_AZ1
Thioflavicoccus_mobilis_8321_8321
Thiolapillus_brandeum_Hiromi_1
Halorhodospira_halochloris_DSM_1059
Halorhodospira_halophila_SL1
Alkalilimnicola_ehrlichii_MLHE_1
Spiribacter_curvatus_UAH_SP71
Spiribacter_salinus_M19_40
Halofilum_ochraceum_XJ16
Oceanococcus_atlanticus_22II_S10r2
Thioalkalivibrio_thiocyanoxidans_ARh2
Thioalkalivibrio_versutus_D301
Thioalkalivibrio_halophilus_HL17
Thioalkalivibrio_nitratireducens_DSM_14787
Thioalkalivibrio_paradoxus_ARh_1
Thioalkalivibrio_denitrificans_ALJD
Thioalkalivibrio_sulfidiphilus_HL_EbGr7
Nitrosococcus_oceani_ATCC_19707
Nitrosococcus_watsonii_C_113
Nitrosococcus_halophilus_Nc_4
Immundisolibacter_cernigliae_TR32
Methylobacterium_aquaticum_MA_22A
Methylobacterium_nodulans_ORS_2060
Methylobacterium_gossipiicola_Gh_105
Methylobacterium_phyllostachyos_BL47
Limnobacter_sp_MED105
Propionivibrio_dicarboxylicus_DSM_5885
0.1

Figure S4

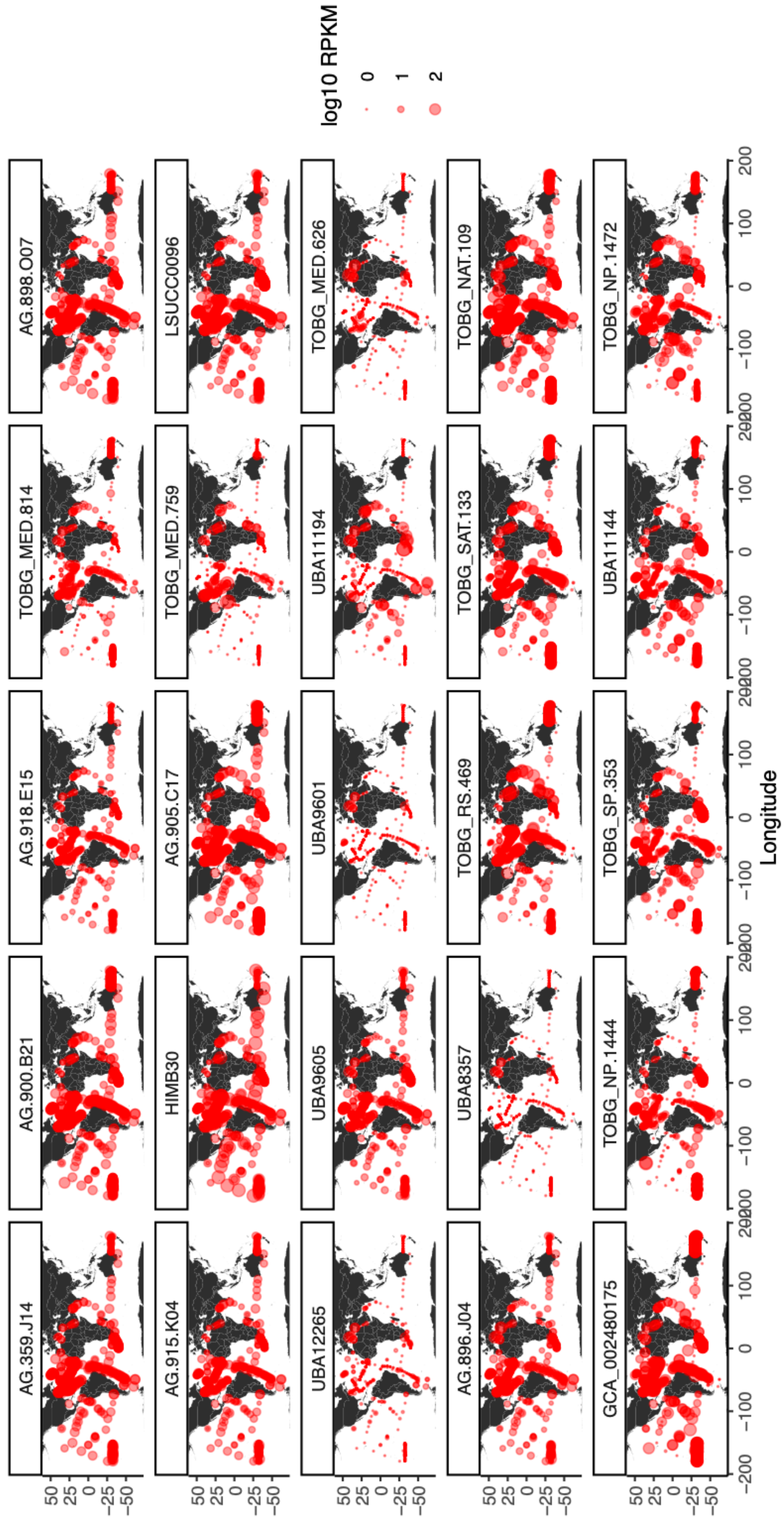

log10 RPKM
0
1
2
AG.898.O07
TOBG_MED.814
AG.918.E15
AG.900.B21
AG.359.J14
LSUCC0096
TOBG_MED.759
AG.905.C17
HIMB30
AG.915.K04
TOBG_MED.626
UBA11194
UBA9601
UBA9605
UBA12265
TOBG_NAT.109
TOBG_SAT.133
TOBG_RS.469
UBA8357
AG.896.J04
TOBG_NP.1472
UBA11144
TOBG_SP.353
TOBG_NP.1444
GCA_002480175
Latitude
Longitude

Figure S5

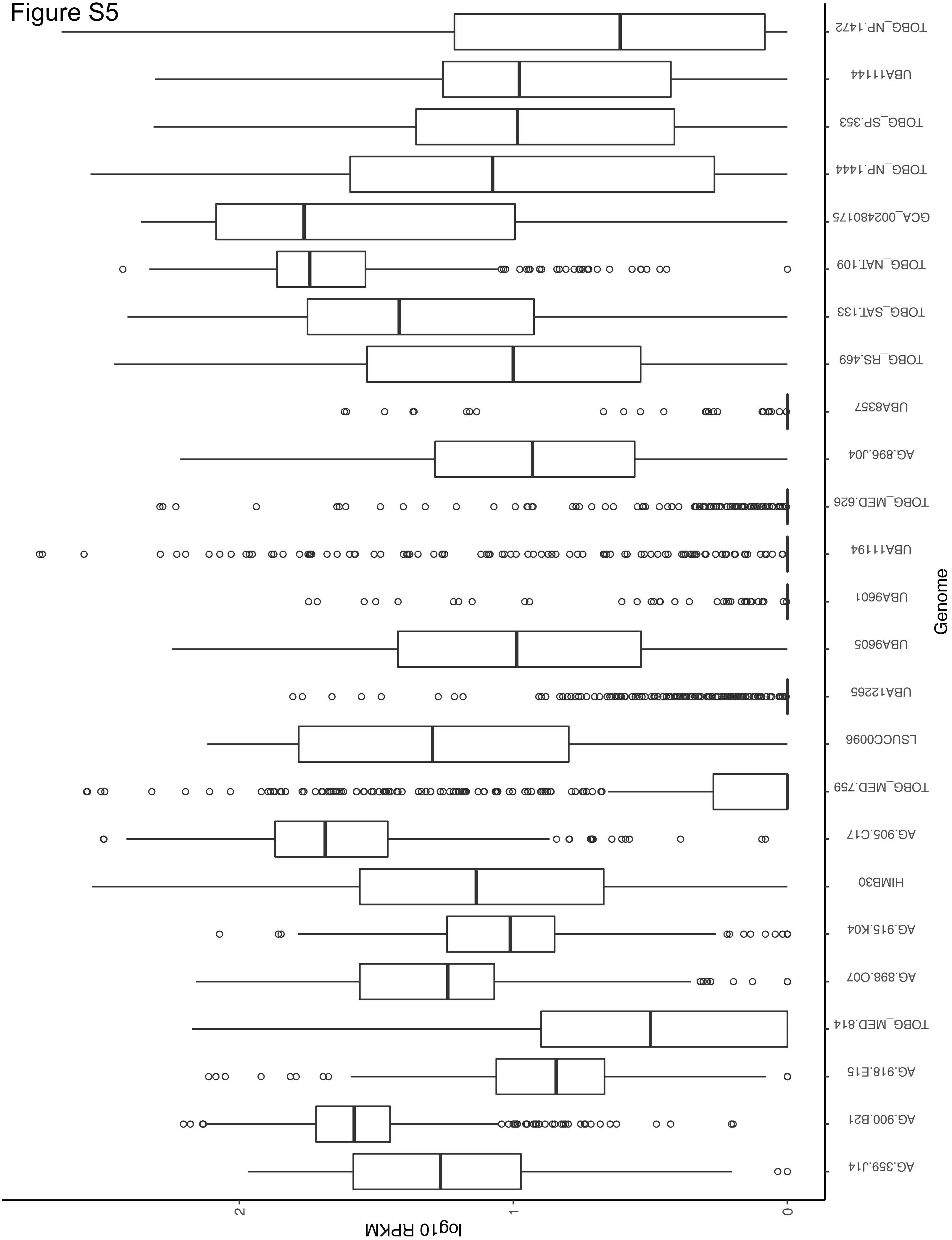

TOBG_NP.1472
UBA11144
TOBG_SP.353
TOBG_NP.1444
GCA_002480175
TOBG_NAT.109
TOBG_SAT.133
TOBG_RS.469
UBA8357
AG.896.J04
TOBG_MED.626
UBA11194
UBA9601
UBA9605
UBA12265
LSUCC0096
TOBG_MED.759
AG.905.C17
HIMB30
AG.915.K04
AG.898.O07
TOBG_MED.814
AG.918.E15
AG.900.B21
AG.359.J14
Genome
2
1
0
log10 RPKM

### Figure S6

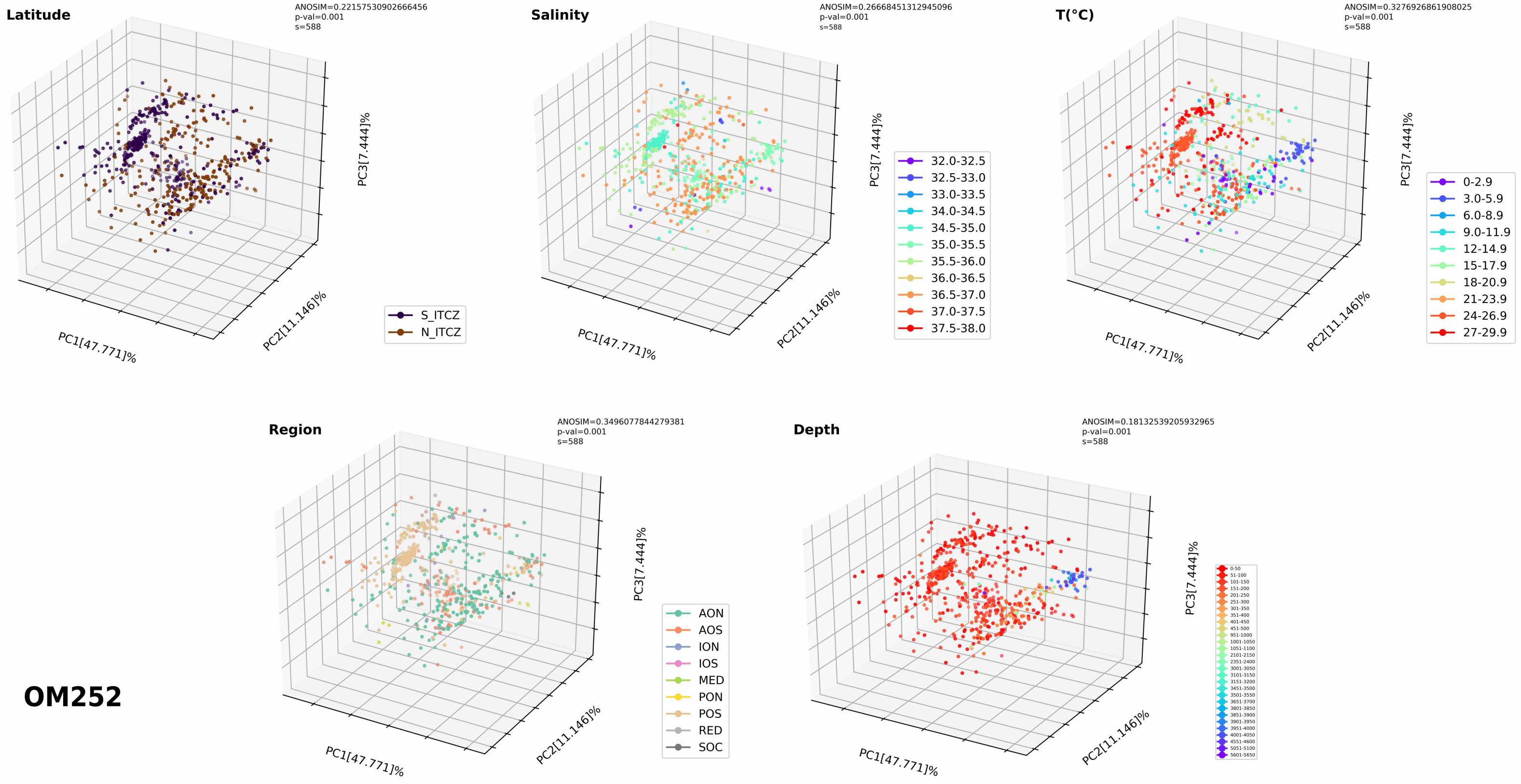

Latitude
ANOSIM=0.22157530902666456
p-val=0.001
s=588
PC1[47.771]%
PC2[11.146]%
PC3[7.444]%
S_ITCZ
N_ITCZ
Salinity
ANOSIM=0.26668451312945096
p-val=0.001
s=588
PC1[47.771]%
PC2[11.146]%
PC3[7.444]%
32.0-32.5
32.5-33.0
33.0-33.5
34.0-34.5
34.5-35.0
35.0-35.5
35.5-36.0
36.0-36.5
36.5-37.0
37.0-37.5
37.5-38.0
T(°C)
ANOSIM=0.3276926861908025
p-val=0.001
s=588
PC1[47.771]%
PC2[11.146]%
PC3[7.444]%
0-2.9
3.0-5.9
6.0-8.9
9.0-11.9
12-14.9
15-17.9
18-20.9
21-23.9
24-26.9
27-29.9
Region
ANOSIM=0.3496077844279381
p-val=0.001
s=588
PC1[47.771]%
PC2[11.146]%
PC3[7.444]%
AON
AOS
ION
IOS
MED
PON
POS
RED
SOC
Depth
ANOSIM=0.18132539205932965
p-val=0.001
s=588
PC1[47.771]%
PC2[11.146]%
PC3[7.444]%
OM252

S7

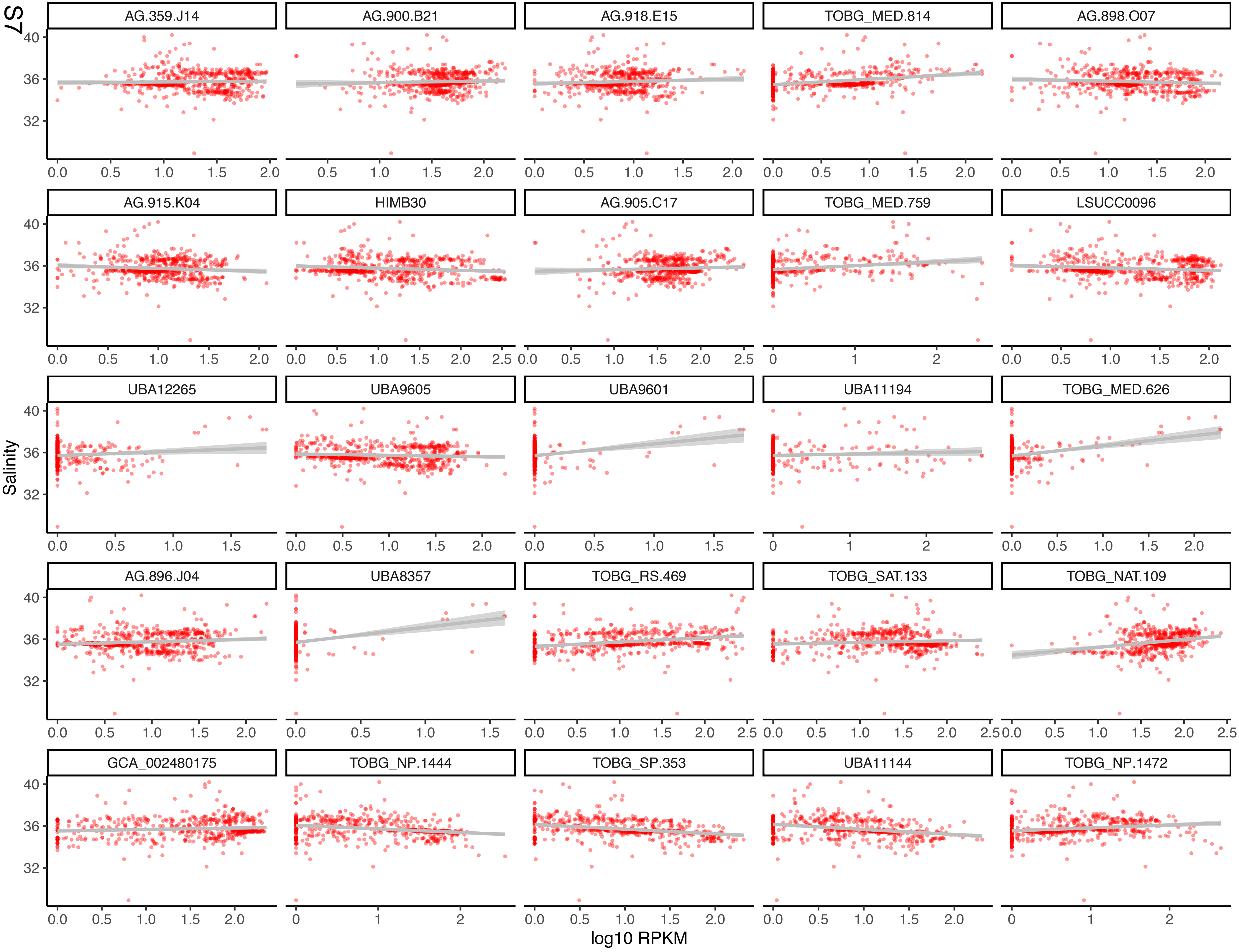
AG.359.J14
AG.900.B21
AG.918.E15
TOBG_MED.814
AG.898.O07
AG.915.K04
HIMB30
AG.905.C17
TOBG_MED.759
LSUCC0096
UBA12265
UBA9605
UBA9601
UBA11194
TOBG_MED.626
AG.896.J04
UBA8357
TOBG_RS.469
TOBG_SAT.133
TOBG_NAT.109
GCA_002480175
TOBG_NP.1444
TOBG_SP.353
UBA11144
TOBG_NP.1472
Salinity
log10 RPKM

S8

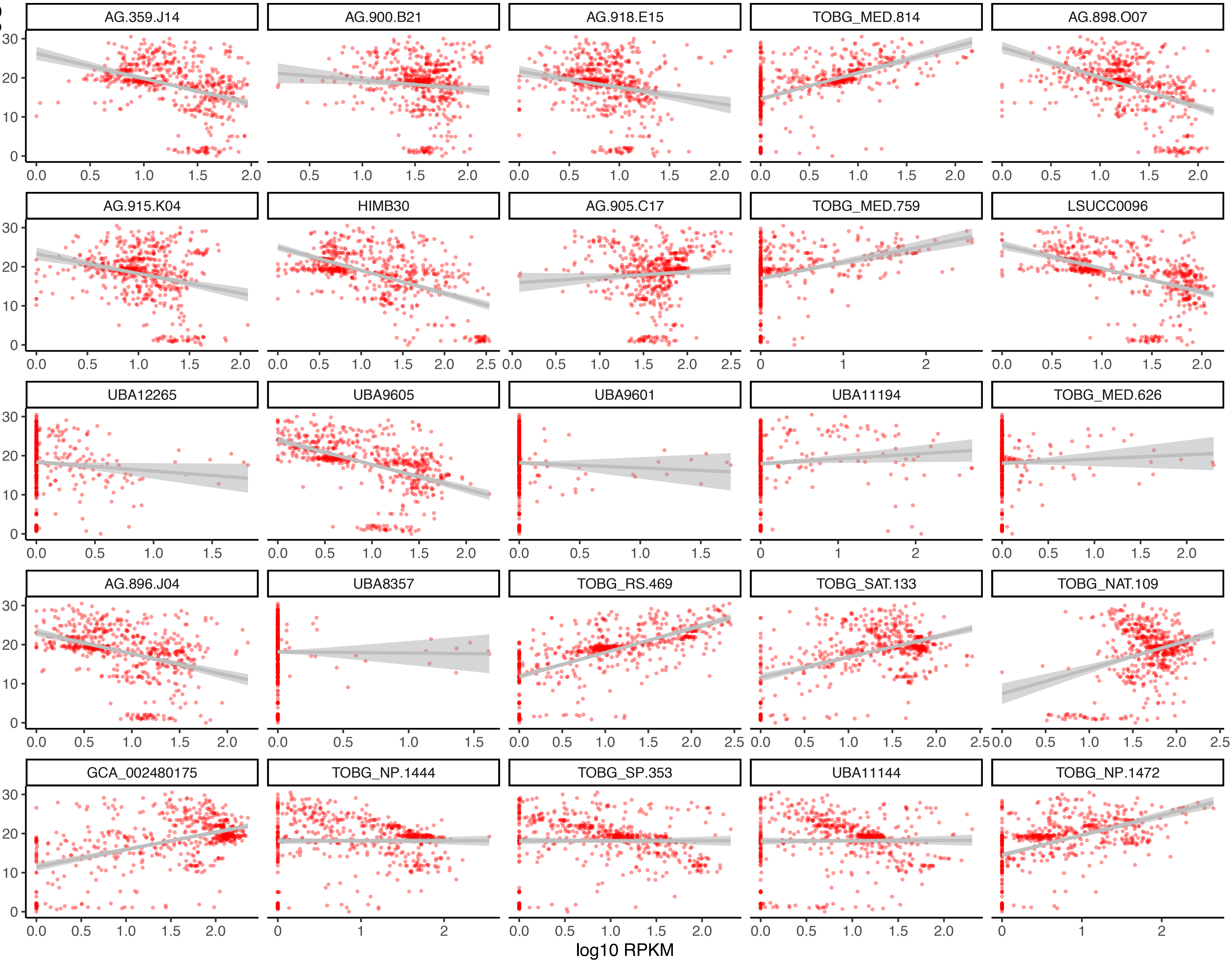
AG.359.J14
AG.900.B21
AG.918.E15
TOBG_MED.814
AG.898.O07
AG.915.K04
HIMB30
AG.905.C17
TOBG_MED.759
LSUCC0096
UBA12265
UBA9605
UBA9601
UBA11194
TOBG_MED.626
AG.896.J04
UBA8357
TOBG_RS.469
TOBG_SAT.133
TOBG_NAT.109
GCA_002480175
TOBG_NP.1444
TOBG_SP.353
UBA11144
TOBG_NP.1472
Temp (..C)
log10 RPKM

S9

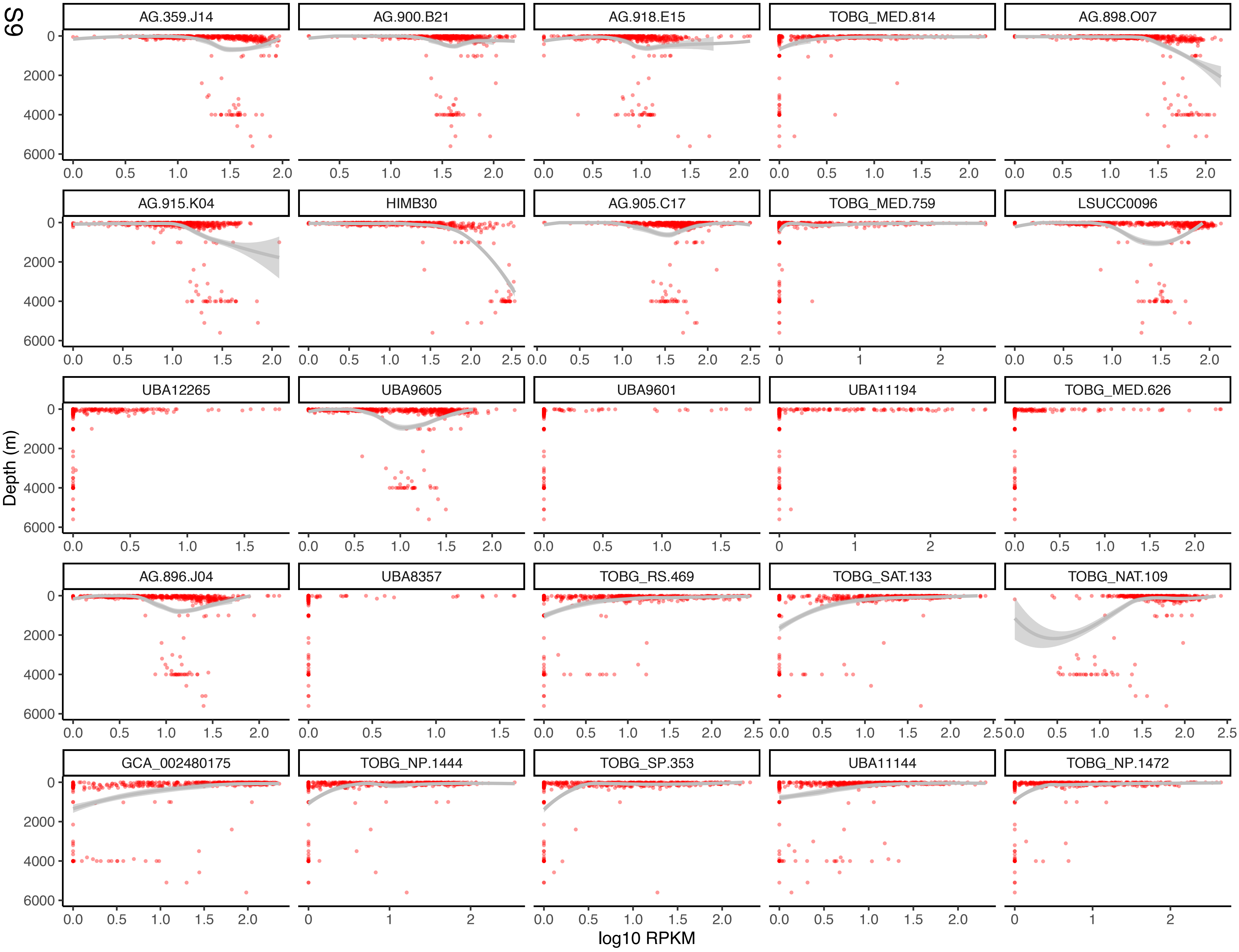
AG.359.J14
AG.900.B21
AG.918.E15
TOBG_MED.814
AG.898.O07
AG.915.K04
HIMB30
AG.905.C17
TOBG_MED.759
LSUCC0096
UBA12265
UBA9605
UBA9601
UBA11194
TOBG_MED.626
AG.896.J04
UBA8357
TOBG_RS.469
TOBG_SAT.133
TOBG_NAT.109
GCA_002480175
TOBG_NP.1444
TOBG_SP.353
UBA11144
TOBG_NP.1472
Depth (m)
log10 RPKM

### Figure S10

A

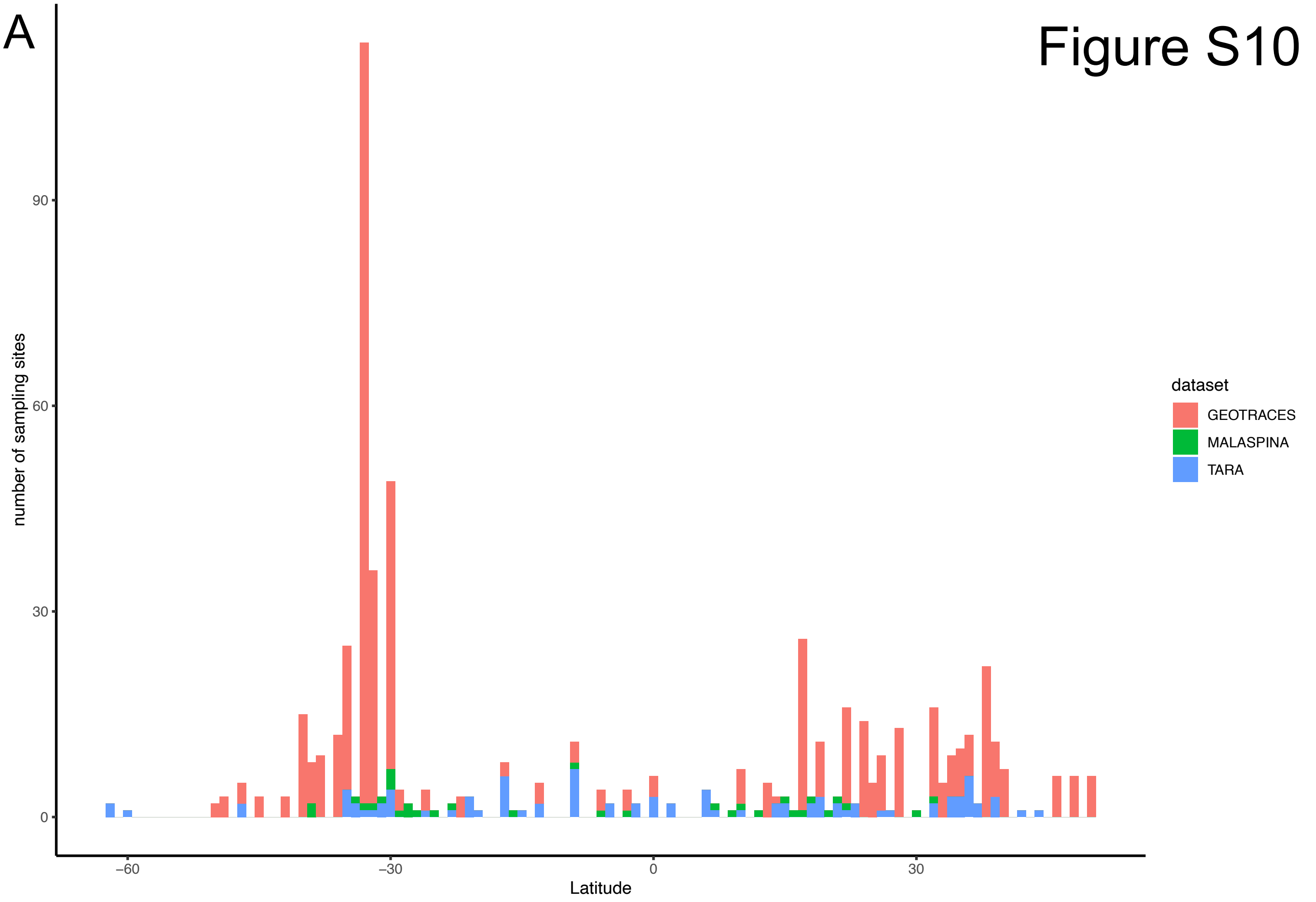


B

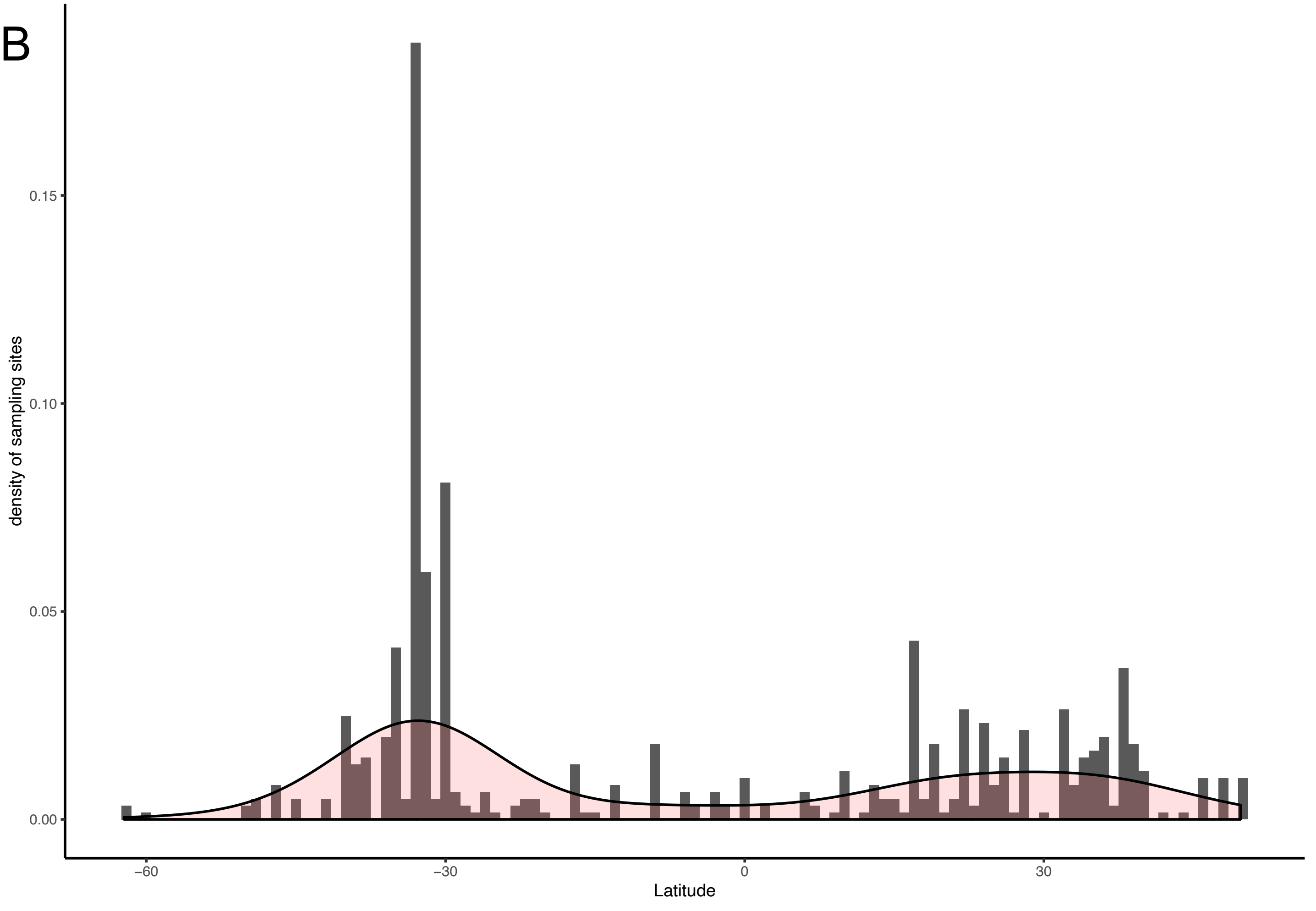

S11

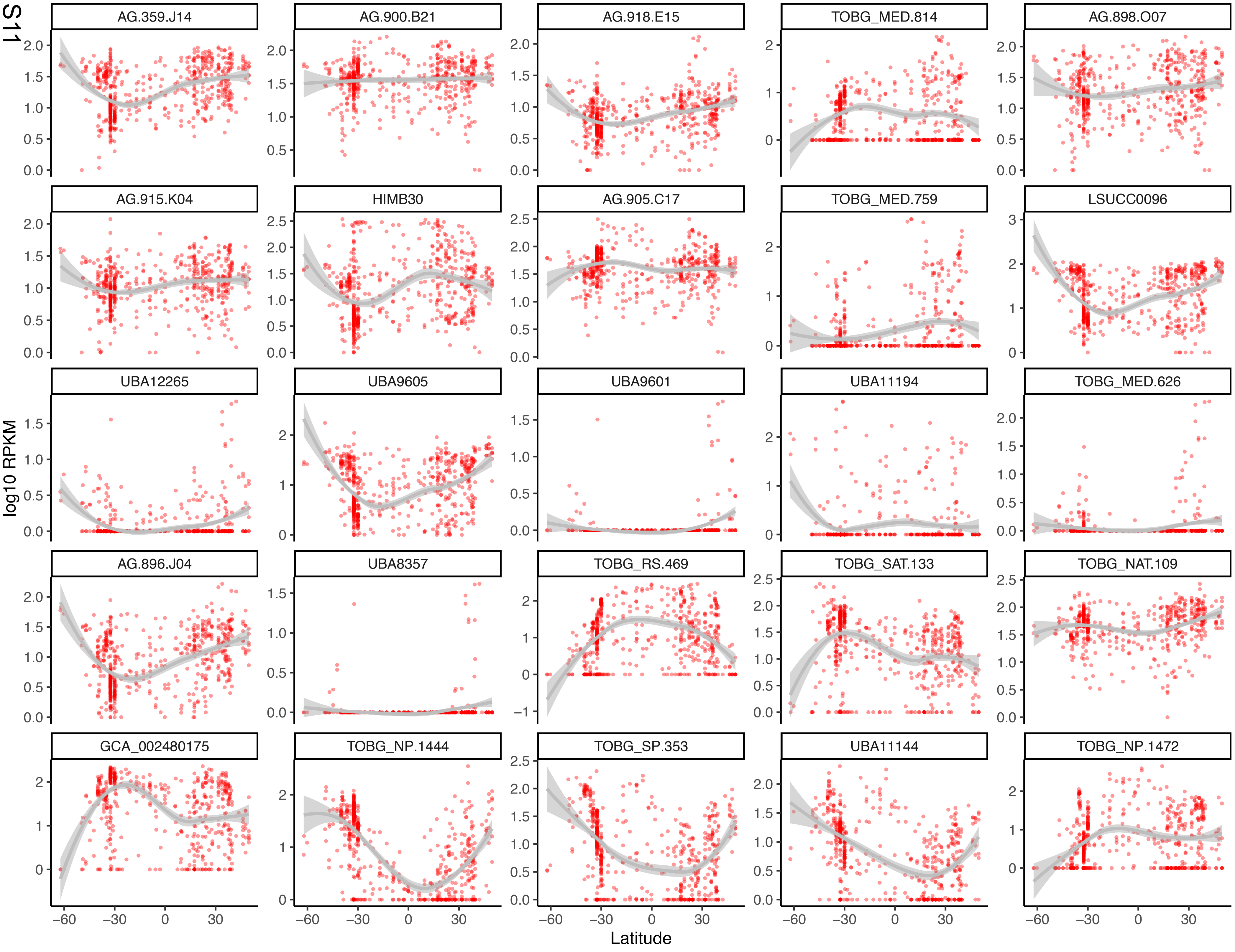
AG.359.J14
AG.900.B21
AG.918.E15
TOBG_MED.814
AG.898.O07
AG.915.K04
HIMB30
AG.905.C17
TOBG_MED.759
LSUCC0096
UBA12265
UBA9605
UBA9601
UBA11194
TOBG_MED.626
AG.896.J04
UBA8357
TOBG_RS.469
TOBG_SAT.133
TOBG_NAT.109
GCA_002480175
TOBG_NP.1444
TOBG_SP.353
UBA11144
TOBG_NP.1472
log10 RPKM
Latitude

### Figure S12

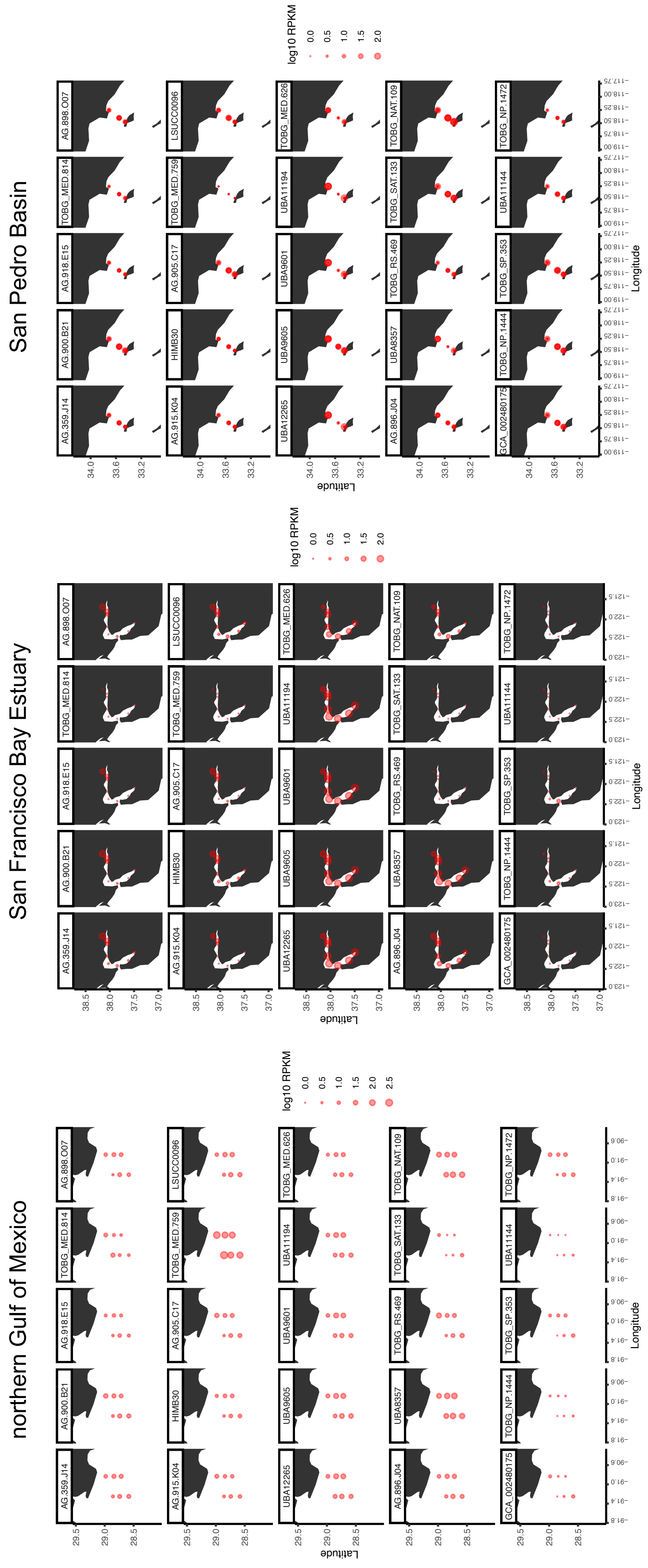
San Pedro Basin
San Francisco Bay Estuary
northern Gulf of Mexico
log10 RPKM
Latitude
Longitude
AG.359.J14
AG.900.B21
AG.918.E15
TOBG_MED.814
AG.898.O07
AG.915.K04
HIMB30
AG.905.C17
TOBG_MED.759
LSUCC0096
UBA12265
UBA9605
UBA9601
UBA11194
TOBG_MED.626
AG.896.J04
UBA8357
TOBG_RS.469
TOBG_SAT.133
TOBG_NAT.109
GCA_002480175
TOBG_NP.1444
TOBG_SP.353
UBA11144
TOBG_NP.1472

Figure S13

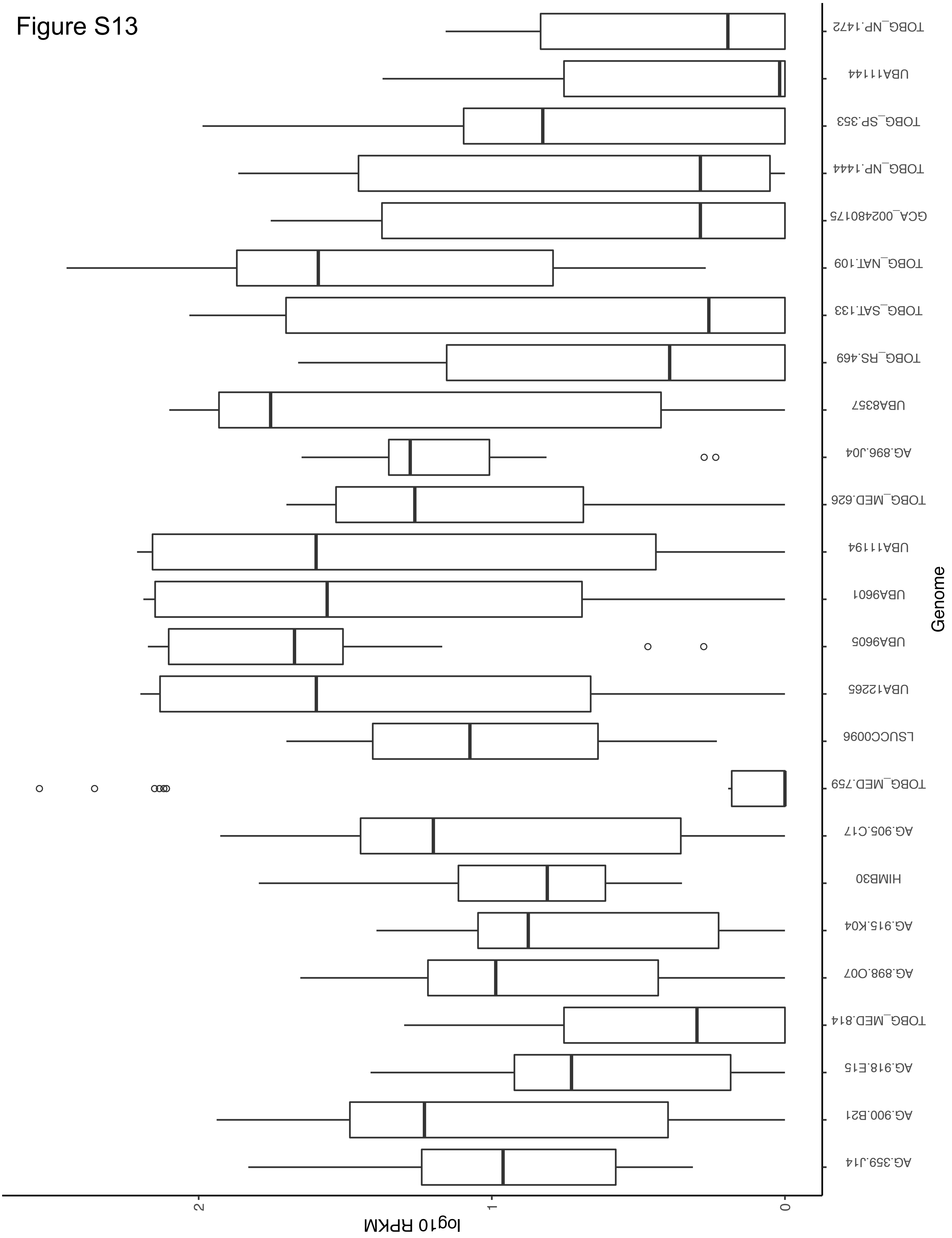

Genome
log10 RPKM
0
1
2
TOBG_NP.1472
UBA11144
TOBG_SP.353
TOBG_NP.1444
GCA_002480175
TOBG_NAT.109
TOBG_SAT.133
TOBG_RS.469
UBA8357
AG.896.J04
TOBG_MED.626
UBA11194
UBA9601
UBA9605
UBA12265
LSUCC0096
TOBG_MED.759
AG.905.C17
HIMB30
AG.915.K04
AG.898.O07
TOBG_MED.814
AG.918.E15
AG.900.B21
AG.359.J14

Figure S14

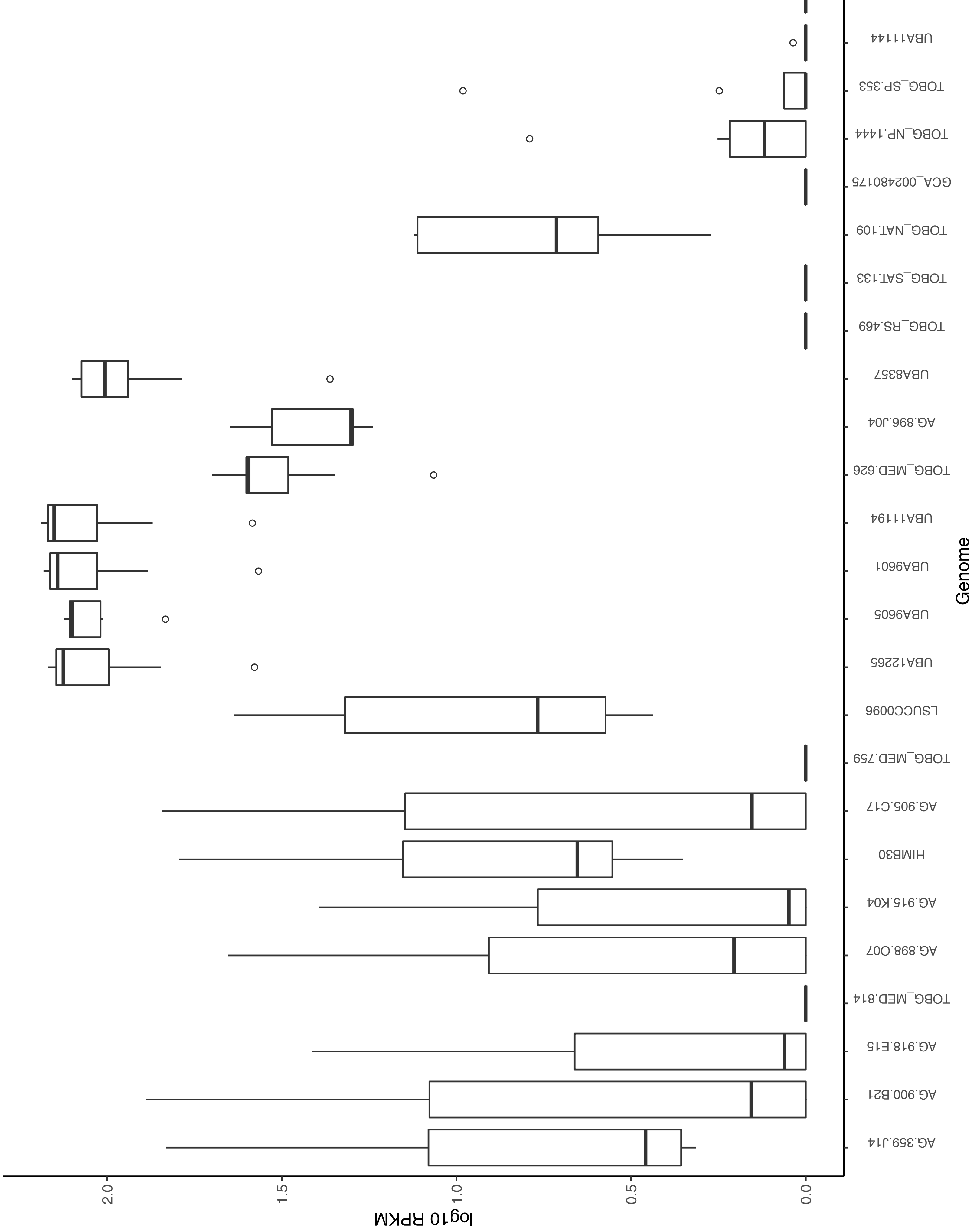
Genome
TOBG_NP.1472
UBA11144
TOBG_SP.353
TOBG_NP.1444
GCA_002480175
TOBG_NAT.109
TOBG_SAT.133
TOBG_RS.469
UBA8357
AG.896.J04
TOBG_MED.626
UBA11194
UBA9601
UBA9605
UBA12265
LSUCC0096
TOBG_MED.759
AG.905.C17
HIMB30
AG.915.K04
AG.898.O07
TOBG_MED.814
AG.918.E15
AG.900.B21
AG.359.J14
log10 RPKM
0.0
0.5
1.0
1.5
2.0

S15

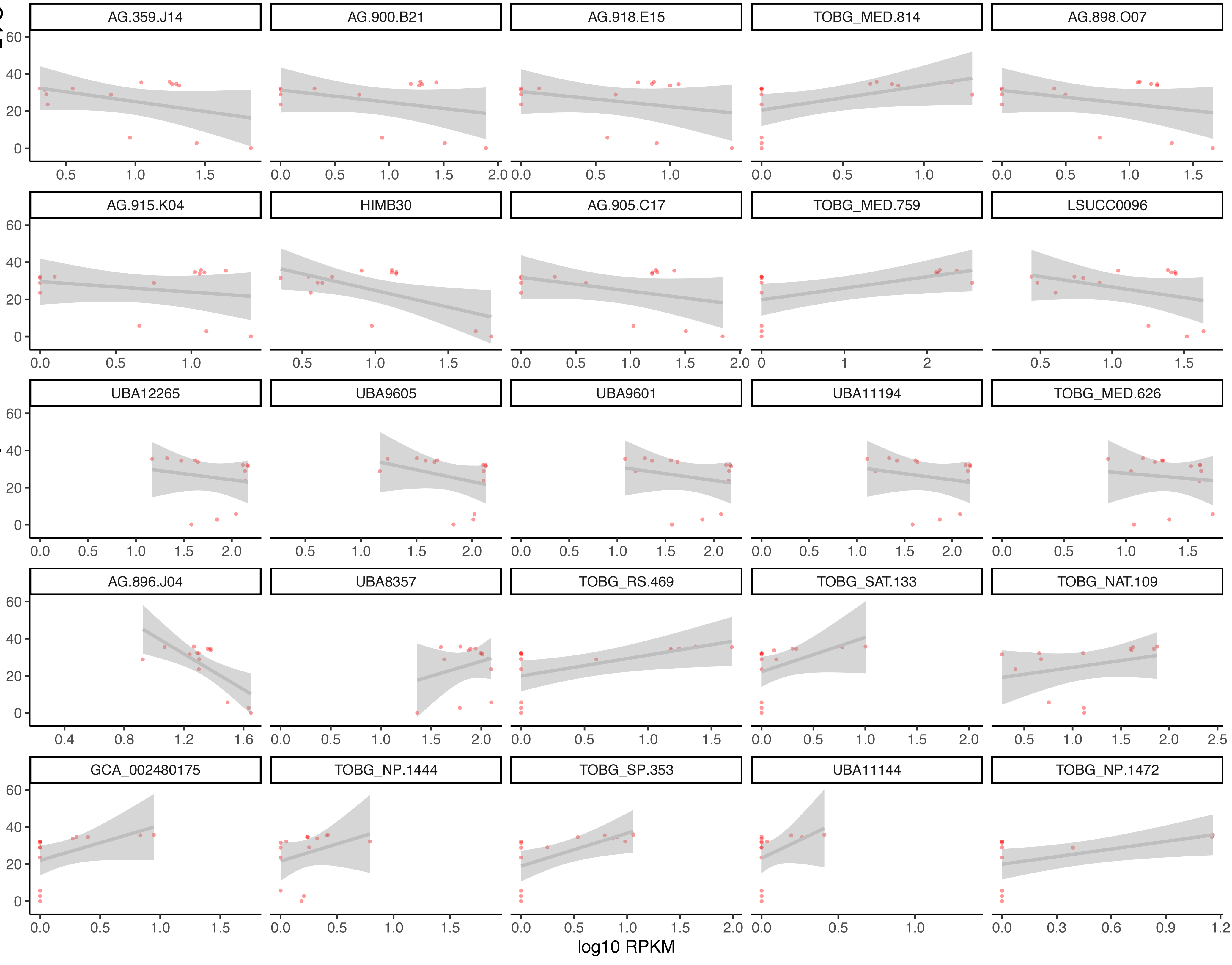

AG.359.J14
AG.900.B21
AG.918.E15
TOBG_MED.814
AG.898.O07
AG.915.K04
HIMB30
AG.905.C17
TOBG_MED.759
LSUCC0096
UBA12265
UBA9605
UBA9601
UBA11194
TOBG_MED.626
AG.896.J04
UBA8357
TOBG_RS.469
TOBG_SAT.133
TOBG_NAT.109
GCA_002480175
TOBG_NP.1444
TOBG_SP.353
UBA11144
TOBG_NP.1472
Salinity
log10 RPKM

Figure S16

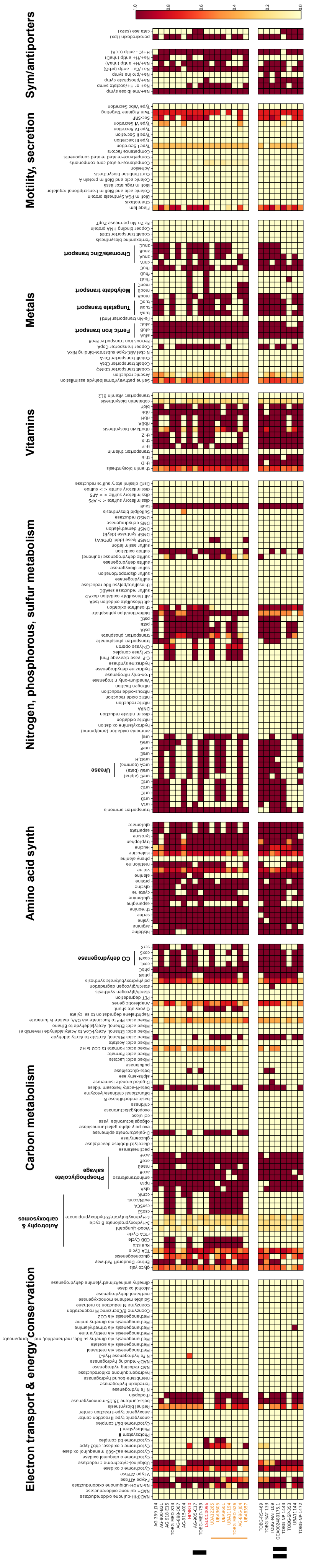
Sym/antiporters
Motility, secretion
Metals
Vitamins
Nitrogen, phosphorous, sulfur metabolism
Amino acid synth
Carbon metabolism
Electron transport & energy conservation
I
II

Figure S17

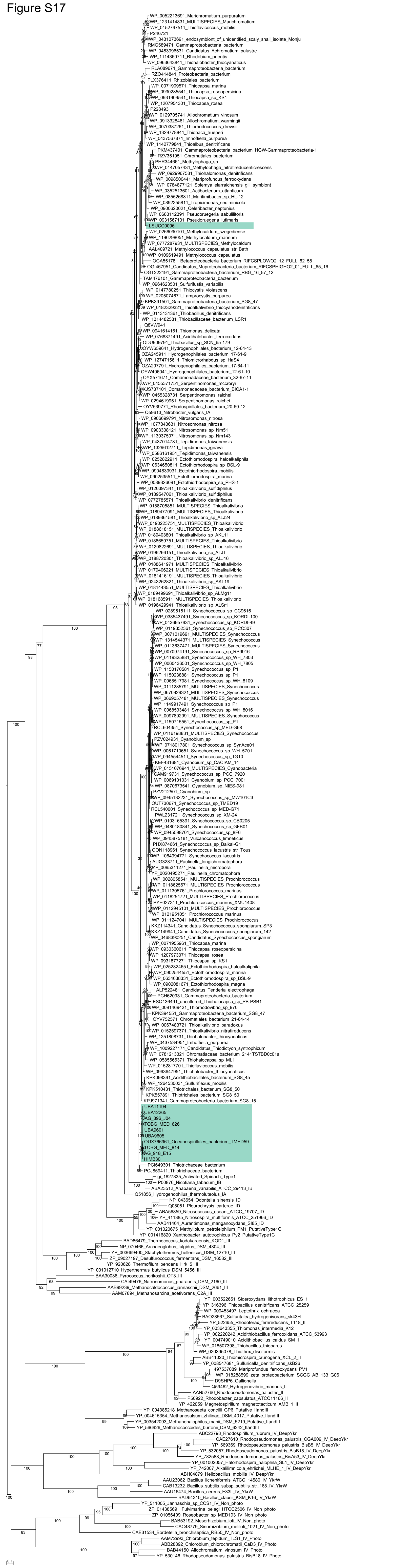
WP_0052213691_Marichromatium_purpuratum
WP_1231414831_MULTISPECIES_Marichromatium
WP_0152797511_Thioflavicoccus_mobilis
P246721
WP_0431073691_endosymbiont_of_unidentified_scaly_snail_isolate_Monju
RMG589471_Gammaproteobacteria_bacterium
WP_0483996531_Candidatus_Achromatium_palustre
WP_1114360711_Rhodobium_orientis
WP_0963643841_Thiohalobacter_thiocyanaticus
RLA089671_Gammaproteobacteria_bacterium
RZO414841_Proteobacteria_bacterium
PLX376411_Rhizobiales_bacterium
WP_0071909571_Thiocapsa_marina
WP_0930285541_Thiocapsa_roseopersicina
WP_0931909541_Thiocapsa_sp_KS1
WP_1207954301_Thiocapsa_rosea
P228493
WP_0129705741_Allochromatium_vinosum
WP_0913328461_Allochromatium_warmingii
WP_0070387261_Thiorhodococcus_drewsii
WP_1329778841_Thiobaca_trueperi
WP_0437567871_Imhoffiella_purpurea
WP_1142779841_Thioalbus_denitrificans
PKM437401_Gammaproteobacteria_bacterium_HGW-Gammaproteobacteria-1
RZV351951_Chromatiales_bacterium
PHR344661_Methylophaga_sp
WP_0147057431_Methylophaga_nitratireducenticrescens
WP_0929967581_Thiohalomonas_denitrificans
WP_0098500441_Mariprofundus_ferrooxydans
WP_0784877121_Solemya_elarraichensis_gill_symbiont
WP_0352513601_Actibacterium_atlanticum
WP_0855268811_Maritimibacter_sp_HL-12
WP_0892355811_Tropicimonas_sediminicola
WP_0900620021_Celeribacter_neptunius
WP_0683112391_Pseudoruegeria_sabulilitoris
WP_0931567131_Pseudoruegeria_lutimaris
LSUCC0096
WP_0266090101_Methylocaldum_szegediense
WP_1196298051_Methylocaldum_marinum
WP_0777287931_MULTISPECIES_Methylocaldum
AAL409721_Methylococcus_capsulatus_str_Bath
WP_0109619491_Methylococcus_capsulatus
OGA551781_Betaproteobacteria_bacterium_RIFCSPLOWO2_12_FULL_62_58
OGI467951_Candidatus_Muproteobacteria_bacterium_RIFCSPHIGHO2_01_FULL_65_16
OGT222191_Gammaproteobacteria_bacterium_RBG_16_57_12
TAM476101_Gammaproteobacteria_bacterium
WP_0964623501_Sulfurifustis_variabilis
WP_0147780251_Thiocystis_violascens
WP_0205074671_Lamprocystis_purpurea
KPK391501_Gammaproteobacteria_bacterium_SG8_47
WP_0183293211_Thioalkalivibrio_thiocyanodenitrificans
WP_0113131361_Thiobacillus_denitrificans
WP_1314482581_Thiobacillaceae_bacterium_LSR1
Q8VW941
WP_0941614161_Thiomonas_delicata
WP_0768371491_Acidihalobacter_ferrooxidans
ODU909791_Thiobacillus_sp_SCN_65-179
OYW659641_Hydrogenophilales_bacterium_12-64-13
OZA245911_Hydrogenophilales_bacterium_17-61-9
WP_1274715611_Thiomicrorhabdus_sp_HaS4
OZA297791_Hydrogenophilales_bacterium_17-64-11
OYW406041_Hydrogenophilales_bacterium_12-61-10
OYX571671_Comamonadaceae_bacterium_32-67-11
WP_0455371751_Serpentinomonas_mccroryi
KJS737101_Comamonadaceae_bacterium_BICA1-1
WP_0455328731_Serpentinomonas_raichei
WP_0294619951_Serpentinomonas_raichei
OYV539771_Rhodospirillales_bacterium_20-60-12
Q59613_Nitrobacter_vulgaris_IA
WP_0906699791_Nitrosomonas_nitrosa
WP_1077843631_Nitrosomonas_nitrosa
WP_0903308121_Nitrosomonas_sp_Nm51
WP_1130375071_Nitrosomonas_sp_Nm143
WP_0437014781_Tepidimonas_taiwanensis
WP_1329612711_Tepidimonas_ignava
WP_0586161951_Tepidimonas_taiwanensis
WP_0252822911_Ectothiorhodospira_haloalkaliphila
WP_0634650811_Ectothiorhodospira_sp_BSL-9
WP_0904839931_Ectothiorhodospira_mobilis
WP_0902535511_Ectothiorhodospira_marina
WP_0089326091_Ectothiorhodospira_sp_PHS-1
WP_0126397341_Thioalkalivibrio_sulfidiphilus
WP_0189547061_Thioalkalivibrio_sulfidiphilus
WP_0772785571_Thioalkalivibrio_denitrificans
WP_0188705851_MULTISPECIES_Thioalkalivibrio
WP_0189477091_MULTISPECIES_Thioalkalivibrio
WP_0189361581_Thioalkalivibrio_sp_ALJ24
WP_0190223751_MULTISPECIES_Thioalkalivibrio
WP_0188618151_MULTISPECIES_Thioalkalivibrio
WP_0189403801_Thioalkalivibrio_sp_AKL11
WP_0188659751_MULTISPECIES_Thioalkalivibrio
WP_0129822691_MULTISPECIES_Thioalkalivibrio
WP_0196266151_Thioalkalivibrio_sp_ALJT
WP_0188720301_Thioalkalivibrio_sp_ALJ16
WP_0188641971_MULTISPECIES_Thioalkalivibrio
WP_0179406221_MULTISPECIES_Thioalkalivibrio
WP_0181416191_MULTISPECIES_Thioalkalivibrio
WP_0243262821_Thioalkalivibrio_sp_AKL19
WP_0181443551_MULTISPECIES_Thioalkalivibrio
WP_0189499691_Thioalkalivibrio_sp_ALMg11
WP_0181685911_MULTISPECIES_Thioalkalivibrio
WP_0196429941_Thioalkalivibrio_sp_ALSr1
WP_0289515111_Synechococcus_sp_CC9616
WP_0385437491_Synechococcus_sp_KORDI-100
WP_0436957931_Synechococcus_sp_KORDI-49
WP_0119352361_Synechococcus_sp_RCC307
WP_0071019691_MULTISPECIES_Synechococcus
WP_1314544371_MULTISPECIES_Synechococcus
WP_0113637471_MULTISPECIES_Synechococcus
WP_0070974191_Synechococcus_sp_RS9916
WP_0119325881_Synechococcus_sp_WH_7803
WP_0060436501_Synechococcus_sp_WH_7805
WP_1150170581_Synechococcus_sp_P1
WP_1150238881_Synechococcus_sp_P1
WP_0068517981_Synechococcus_sp_WH_8109
WP_0111285791_MULTISPECIES_Synechococcus
WP_0670929321_MULTISPECIES_Synechococcus
WP_0669057481_MULTISPECIES_Synechococcus
WP_1149917491_Synechococcus_sp_P1
WP_0068533481_Synechococcus_sp_WH_8016
WP_0097892991_MULTISPECIES_Synechococcus
WP_1150715551_Synechococcus_sp_P1
RCL604351_Synechococcus_sp_MED-G68
WP_0116198831_MULTISPECIES_Synechococcus
PZV024931_Cyanobium_sp
WP_0718017801_Synechococcus_sp_SynAce01
WP_0061710651_Synechococcus_sp_WH_5701
WP_0945544511_Synechococcus_sp_1G10
KEF431681_Cyanobium_sp_CACIAM_14
WP_0151076941_MULTISPECIES_Cyanobacteria
CAM919731_Synechococcus_sp_PCC_7920
WP_0069101031_Cyanobium_sp_PCC_7001
WP_0870673541_Cyanobium_sp_NIES-981
PZV212501_Cyanobium_sp
WP_0945132231_Synechococcus_sp_MW101C3
OUT730671_Synechococcus_sp_TMED19
RCL540001_Synechococcus_sp_MED-G71
PWL231721_Synechococcus_sp_XM-24
WP_0103165391_Synechococcus_sp_CB0205
WP_0480180841_Synechococcus_sp_GFB01
WP_0945598701_Synechococcus_sp_8F6
WP_0945875181_Vulcanococcus_limneticus
PHX874661_Synechococcus_sp_Baikal-G1
OON118961_Synechococcus_lacustris_str_Tous
WP_1064994771_Synechococcus_lacustris
AUG328711_Paulinella_longichromatophora
YP_0095311271_Paulinella_micropora
YP_0020495271_Paulinella_chromatophora
WP_0028058541_MULTISPECIES_Prochlorococcus
WP_0118625671_MULTISPECIES_Prochlorococcus
WP_0111305761_Prochlorococcus_marinus
WP_0118254721_MULTISPECIES_Prochlorococcus
PYE027311_Prochlorococcus_marinus_XMU1408
WP_0112945101_MULTISPECIES_Prochlorococcus
WP_0121951051_Prochlorococcus_marinus
WP_0111247041_MULTISPECIES_Prochlorococcus
KKZ114341_Candidatus_Synechococcus_spongiarum_SP3
KKZ149941_Candidatus_Synechococcus_spongiarum_142
WP_0468390251_Candidatus_Synechococcus_spongiarum
WP_0071955961_Thiocapsa_marina
WP_0930360611_Thiocapsa_roseopersicina
WP_1207973071_Thiocapsa_rosea
WP_0931877271_Thiocapsa_sp_KS1
WP_0252824651_Ectothiorhodospira_haloalkaliphila
WP_0902544551_Ectothiorhodospira_marina
WP_0634638331_Ectothiorhodospira_sp_BSL-9
WP_0902081671_Ectothiorhodospira_magna
ALP522481_Candidatus_Tenderia_electrophaga
PCH620931_Gammaproteobacteria_bacterium
ESQ136491_uncultured_Thiohalocapsa_sp_PB-PSB1
WP_0091469421_Thiorhodovibrio_sp_970
KPK394551_Gammaproteobacteria_bacterium_SG8_47
OYV752571_Chromatiales_bacterium_21-64-14
WP_0067483721_Thioalkalivibrio_paradoxus
WP_0152597371_Thioalkalivibrio_nitratireducens
WP_1251808731_Thiohalobacter_thiocyanaticus
WP_0437534951_Imhoffiella_purpurea
WP_1009227171_Candidatus_Thiodictyon_syntrophicum
WP_0781213321_Chromatiaceae_bacterium_2141TSTBD0c01a
WP_0585565371_Thiohalocapsa_sp_ML1
WP_0152817701_Thioflavicoccus_mobilis
WP_0963647951_Thiohalobacter_thiocyanaticus
KPK098391_Acidithiobacillales_bacterium_SG8_45
WP_1264530031_Sulfuriflexus_mobilis
KPK510431_Thiotrichales_bacterium_SG8_50
KPK557891_Thiotrichales_bacterium_SG8_50
KPJ971341_Gammaproteobacteria_bacterium_SG8_15
UBA11194
UBA12265
AG_896_J04
TOBG_MED_626
UBA9601
UBA9605
OUX766961_Oceanospirillales_bacterium_TMED59
TOBG_MED_814
AG_918_E15
HIMB30
PCI649301_Thiotrichaceae_bacterium
PCJ859411_Thiotrichaceae_bacterium
gi_1827835_Activated_Spinach_Type1
P00876_Nicotiana_tabacum_IB
ABA23512_Anabaena_variabilis_ATCC_29413_IB
Q51856_Hydrogenophilus_thermoluteolus_IA
NP_043654_Odontella_sinensis_ID
Q08051_Pleurochrysis_carterae_ID
ABA56859_Nitrosococcus_oceani_ATCC_19707_ID
YP_411385_Nitrosospira_multiformis_ATCC_25196_ID
AAB41464_Aurantimonas_manganoxydans_SI85_ID
YP_001020675_Methylibium_petroleiphilum_PM1_PutativeType1C
YP_001416820_Xanthobacter_autotrophicus_Py2_PutativeType1C
BAD86479_Thermococcus_kodakaraensis_KOD1_III
NP_070466_Archaeoglobus_fulgidus_DSM_4304_III
YP_003669400_Staphylothermus_hellenicus_DSM_12710_III
ZP_09027197_Desulfurococcus_fermentans_DSM_16532_III
YP_920628_Thermofilum_pendens_Hrk_5_III
YP_001012710_Hyperthermus_butylicus_DSM_5456_III
BAA30036_Pyrococcus_horikoshii_OT3_III
CAI49476_Natronomonas_pharaonis_DSM_2160_III
AAB99239_Methanocaldococcus_jannaschii_DSM_2661_III
AAM07894_Methanosarcina_acetivorans_C2A_III
YP_003522651_Sideroxydans_lithotrophicus_ES_1
YP_316396_Thiobacillus_denitrificans_ATCC_25259
WP_009453497_Leptothrix_ochracea
BAO28567_Sulfuritalea_hydrogenivorans_sk43H
YP_522655_Rhodoferax_ferrireducens_T118_II
YP_003643355_Thiomonas_intermedia_K12
WP_002220242_Acidithiobacillus_ferrooxidans_ATCC_53993
WP_004749010_Acidithiobacillus_caldus_SM_1
WP_018507398_Thiobacillus_thioparus
WP_020395078_Thiothrix_disciformis
ABB41020_Thiomicrospira_crunogena_XCL_2_II
YP_008547681_Sulfuricella_denitrificans_skB26
497537089_Mariprofundus_ferrooxydans_PV1
WP_018288599_zeta_proteobacterium_SCGC_AB_133_G06
D9SHP6_Gallionella
Q59462_Hydrogenovibrio_marinus_II
AAN52766_Rhodopseudomonas_palustris_II
P50922_Rhodobacter_capsulatus_ATCC11166_II
YP_422059_Magnetospirillum_magnetotacticum_AMB_1_II
YP_004385218_Methanosaeta_concilii_GP6_Putative_IIandIII
YP_004615354_Methanosalsum_zhilinae_DSM_4017_Putative_IIandIII
YP_003542093_Methanohalophilus_mahii_DSM_5219_Putative_IIandIII
YP_566926_Methanococcoides_burtonii_DSM_6242_IIandIII
ABC22798_Rhodospirillum_rubrum_IV_DeepYkr
CAE27610_Rhodopseudomonas_palustris_CGA009_IV_DeepYkr
YP_569369_Rhodopseudomonas_palustris_BisB5_IV_DeepYkr
YP_532057_Rhodopseudomonas_palustris_BisB18_IV_DeepYkr
YP_782588_Rhodopseudomonas_palustris_BisA53_IV_DeepYkr
YP_001002057_Halorhodospira_halophila_SL1_IV_DeepYkr
YP_742007_Alkalilimnicola_ehrlichei_MLHE_1_IV_DeepYkr
ABH04879_Heliobacillus_mobilis_IV_DeepYkr
AAU23062_Bacillus_licheniformis_ATCC_14580_IV_YkrW
CAB13232_Bacillus_subtilis_subsp_subtilis_str_168_IV_YkrW
AAU16474_Bacillus_cereus_E33L_IV_YkrW
BAD64310_Bacillus_clausii_KSM_K16_IV_YkrW
YP_511005_Jannaschia_sp_CCS1_IV_Non_photo
ZP_01438569_Fulvimarina_pelagi_HTCC2506_IV_Non_photo
ZP_01056409_Roseobacter_sp_MED193_IV_Non_photo
BAB53192_Mesorhizobium_loti_IV_Non_photo
CAC48779_Sinorhizobium_meliloti_1021_IV_Non_photo
CAE31534_Bordetella_bronchiseptica_RB50_IV_Non_photo
AAM72993_Chlorobium_tepidum_TLS1_IV_Photo
ABB28892_Chlorobium_chlorochromatii_CaD3_IV_Photo
BAB44150_Allochromatium_vinosum_IV_Photo
YP_530146_Rhodopseudomonas_palustris_BisB18_IV_Photo

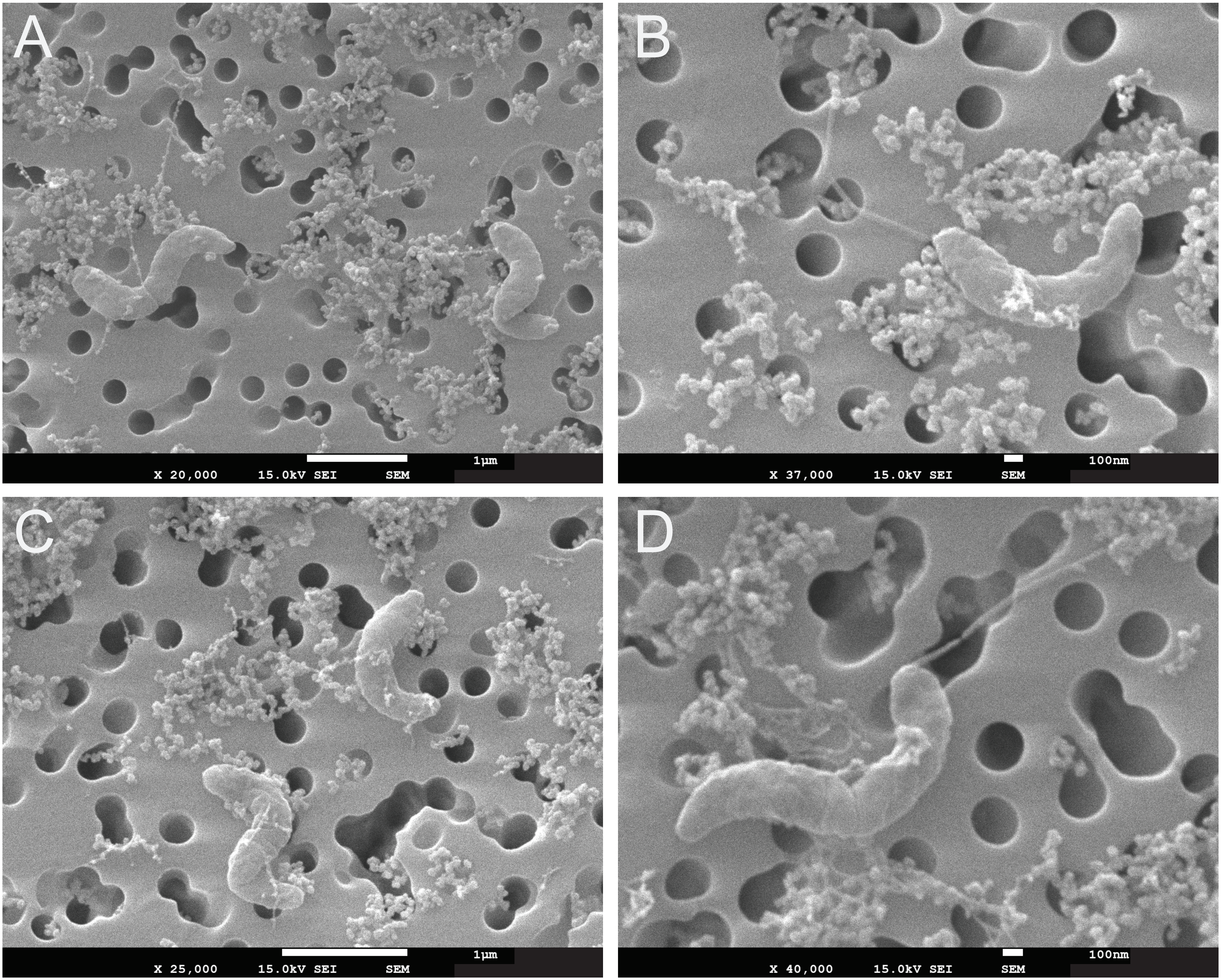
A
1µm
X 20,000 15.0kV SEI SEM
B
100nm
X 37,000 15.0kV SEI SEM
C
1µm
X 25,000 15.0kV SEI SEM
D
100nm
X 40,000 15.0kV SEI SEM

### Figure S19

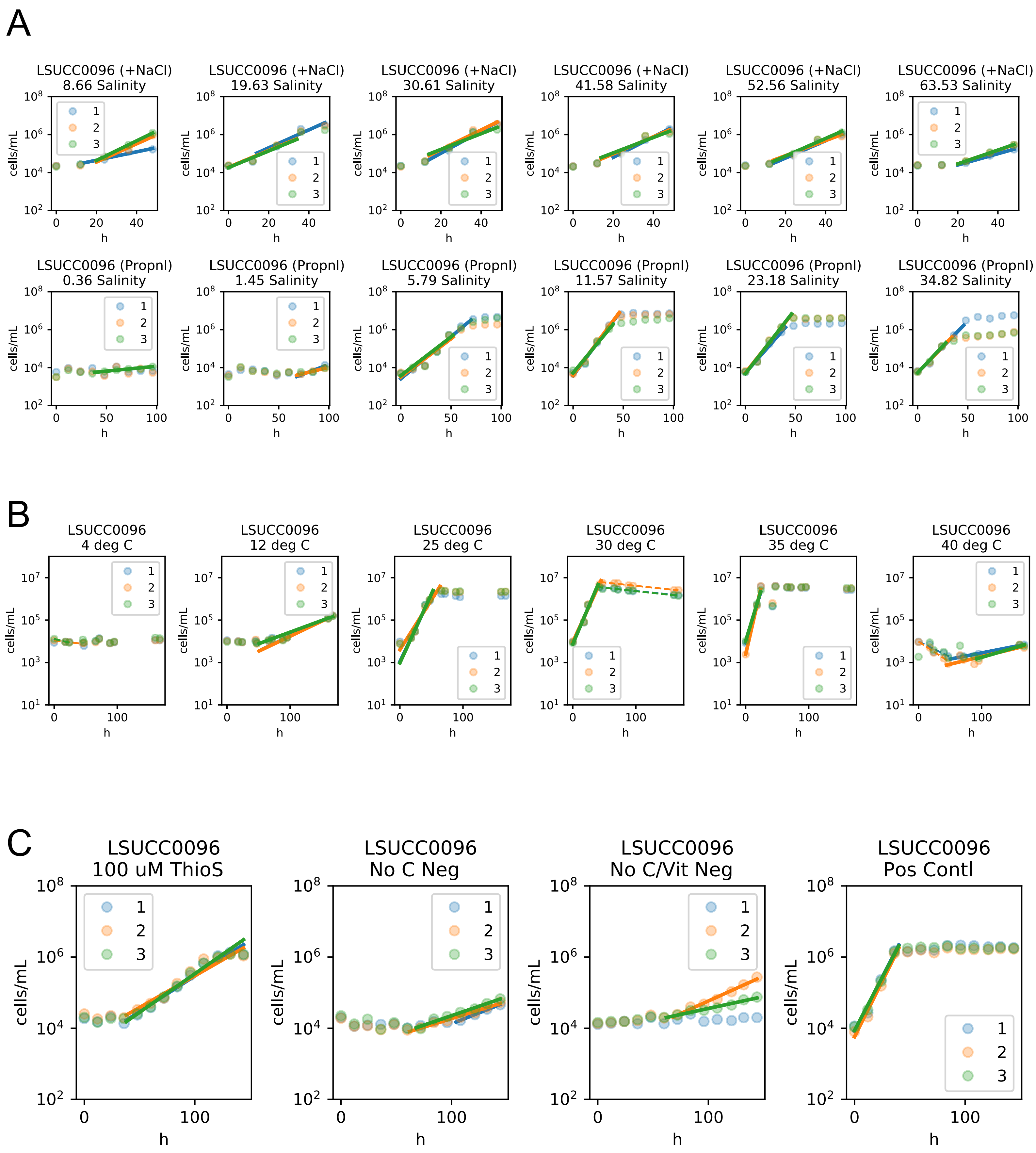
A
LSUCC0096 (+NaCl) 8.66 Salinity
LSUCC0096 (+NaCl) 19.63 Salinity
LSUCC0096 (+NaCl) 30.61 Salinity
LSUCC0096 (+NaCl) 41.58 Salinity
LSUCC0096 (+NaCl) 52.56 Salinity
LSUCC0096 (+NaCl) 63.53 Salinity
LSUCC0096 (Propnl) 0.36 Salinity
LSUCC0096 (Propnl) 1.45 Salinity
LSUCC0096 (Propnl) 5.79 Salinity
LSUCC0096 (Propnl) 11.57 Salinity
LSUCC0096 (Propnl) 23.18 Salinity
LSUCC0096 (Propnl) 34.82 Salinity
B
LSUCC0096 4 deg C
LSUCC0096 12 deg C
LSUCC0096 25 deg C
LSUCC0096 30 deg C
LSUCC0096 35 deg C
LSUCC0096 40 deg C
C
LSUCC0096 100 uM ThioS
LSUCC0096 No C Neg
LSUCC0096 No C/Vit Neg
LSUCC0096 Pos Contl
cells/mL
h
1
2
3
